# Axonal swelling as a neuron-specific compartment to sequester and expel misfolded proteins

**DOI:** 10.64898/2026.05.04.722616

**Authors:** Man Pok Lu, Yingjie Wu, Yi Rong, Yining Chen, Xiaochun Yu, Hongjiang Liu, Minxing Zhang, Yuner Yan, Yuze Chen, Kuankuan Xin, Daniel Schweitzer, Yidong Li, Victor Anggono, Massimo A. Hilliard, Zhaoyu Li

**Affiliations:** Queensland Brain Institute, The University of Queensland, Brisbane, QLD 4072, Australia; Melbourne Brain Centre Imaging Unit, Department of Radiology, The University of Melbourne, Melbourne, VIC 3010, Australia; School of Medicine, The University of Queensland, Brisbane, QLD 4072, Australia; Centre for Neurosciences, Mater Hospital, Brisbane, QLD 4101, Australia

**Keywords:** axonal swelling, aggresome, exopher, microvesicle, SolAS, quality control compartment, SUMO, ubiquitin, SOD1, polydipeptide

## Abstract

Neurons are long-lived, highly specialized cells with extended neurites, requiring precise control of misfolded proteins over time and space. Yet, where misfolded proteins are directed and how quality-control pathways adapt during aging are still unresolved. Here, we identify a neuron-specific quality-control compartment that emerges in neurites as aggresome function declines with age, which we call SolAS (Soluble Misfolded Proteins–induced Axonal Swellings). These structures not only sequester misfolded proteins but also facilitate their clearance via microvesicles and exophers. During aging, neurite SolAS and soma aggresomes function hierarchically to maintain proteostasis, with aggresomes acting as the primary sequestration sites in young neurons and SolAS taking over this role as their function declines with age. This transition is driven by a shift from a ubiquitin-dominant to a SUMO-dominant balance. Moreover, solid pathogenic amyloids, such as GA50, can be converted into soluble forms and sequestered into SolAS via SUMO fusion, thereby reducing neurotoxicity. Our findings identify a previously unrecognized neuronal quality-control pathway critical for proteostasis during aging.

## Introduction

Proteostasis relies on the efficient sequestration and clearance of misfolded proteins through different quality-control pathways. Neurons, as long-lived and highly polarized cells with extended neurites, place significant demands on these quality-control systems. Disruption of proteostasis leads to neural dysfunctions and is associated with many aging-related neurodegenerative diseases^1,2^. However, the mechanisms by which neurons maintain proteostasis during aging remain unclear.

Cells use various mechanisms to process misfolded proteins, including refolding by chaperones and degradation by the proteasome and autophagy^2,3^. When these mechanisms are overwhelmed or inadequate, misfolded proteins are sequestered into specialized quality-control compartments^3^. In yeast, soluble misfolded proteins are directed to JUNQ (Juxtanuclear Quality Control), whereas insoluble misfolded proteins are sequestered into IPODs (Insoluble Protein Deposit)^4^. In mammals, cytosolic misfolded proteins can be sequestered into aggresomes^5,6^. Additionally, misfolded proteins in different cellular organelles can also be directed to specialized compartments, such as the nucleolus for phase-separated nuclear misfolded proteins^7^, the INQ (Intranuclear Quality Control) for nuclear misfolded proteins^8^, and the ERAC (ER-associated compartment) for ER misfolded proteins^9^. However, most of these quality-control compartments are located in the soma. As neurons are highly polarized cells, they must maintain proteostasis not only in somata but also across neurites, raising the question of how misfolded proteins are spatially partitioned and whether neurite-specific sequestration sites exist.

Neurons are exceptionally long-lived cells, with most of them persisting throughout an organism’s lifespan. However, multiple quality-control pathways, including autophagy and proteasome activity, decline with age^10^. How neurons maintain proteostasis when canonical quality-control pathways deteriorate remains poorly understood.

To answer these questions, we introduced misfolded proteins, including Amyotrophic Lateral Sclerosis (ALS/MND)-associated SOD1(A4V) or C9ORF72-derived polyPA and polyGA^11,12^, into *C. elegans* neurons. The short lifespan (∼3 weeks), transparent body, and well-characterized nervous system of this organism make it an excellent system for tracking misfolded proteins in selected neurons throughout their lifespan. By doing this, we identified SolAS (Soluble Misfolded Proteins-induced Axonal Swelling) as a neuron-specific quality-control compartment. Unlike degeneration- or death-associated pathological axonal swellings, SolAS protect neurons by sequestering and expelling soluble misfolded proteins. They function hierarchically with aggresomes in proteostasis. In young animals, aggresomes in somata function as the primary sequestration sites. As aggresome function declines with age, SolAS in neurites emerge as the major reservoir. SolAS also facilitate the removal of misfolded proteins via exophers and microvesicles. The aggresome-SolAS transition is regulated by ubiquitin-SUMO balance, with ubiquitin supporting aggresomes and SUMO promoting SolAS. These findings reveal SolAS as an important mechanism for proteostasis in neurons.

## Results

### Misfolded proteins are sequestered into aggresomes in young *C. elegans* neurons

To investigate how misfolded proteins are processed within neurons, we introduced ALS-associated misfolded proteins SOD1(A4V) and C9ORF72-derived polyPA (PA50)^11,12^ into the *C. elegans* GABAergic motor neurons using *Punc-25* promoter. The cell bodies of these neurons are situated in the ventral cord, with each extending a neurite consisting of a ventral, commissural, and dorsal segment, which allows unambiguous identification (Fig. 1A and S1A; see Methods for detailed information). *C. elegans* develops through four larval stages (L1 to L4) before reaching the adult stage, after which they live for approximately three weeks. At the L4 stage, we observed aggregates in the motor neurons expressing pathogenic proteins SOD1(A4V) or PA50 (Fig. 1B). These aggregates were typically spherical, about 1-2 µm in diameter, located adjacent to the nucleus, and present in approximately 60% of neurons (Fig. 1C). Similar aggregates were also observed when expressing SOD1(A4V) in other neurons, such as the cholinergic motor neurons (*Punc-129* promoter) and the sublateral neurons (*Pacr-15* promoter) (Fig. S1B).

**Fig. 1:**
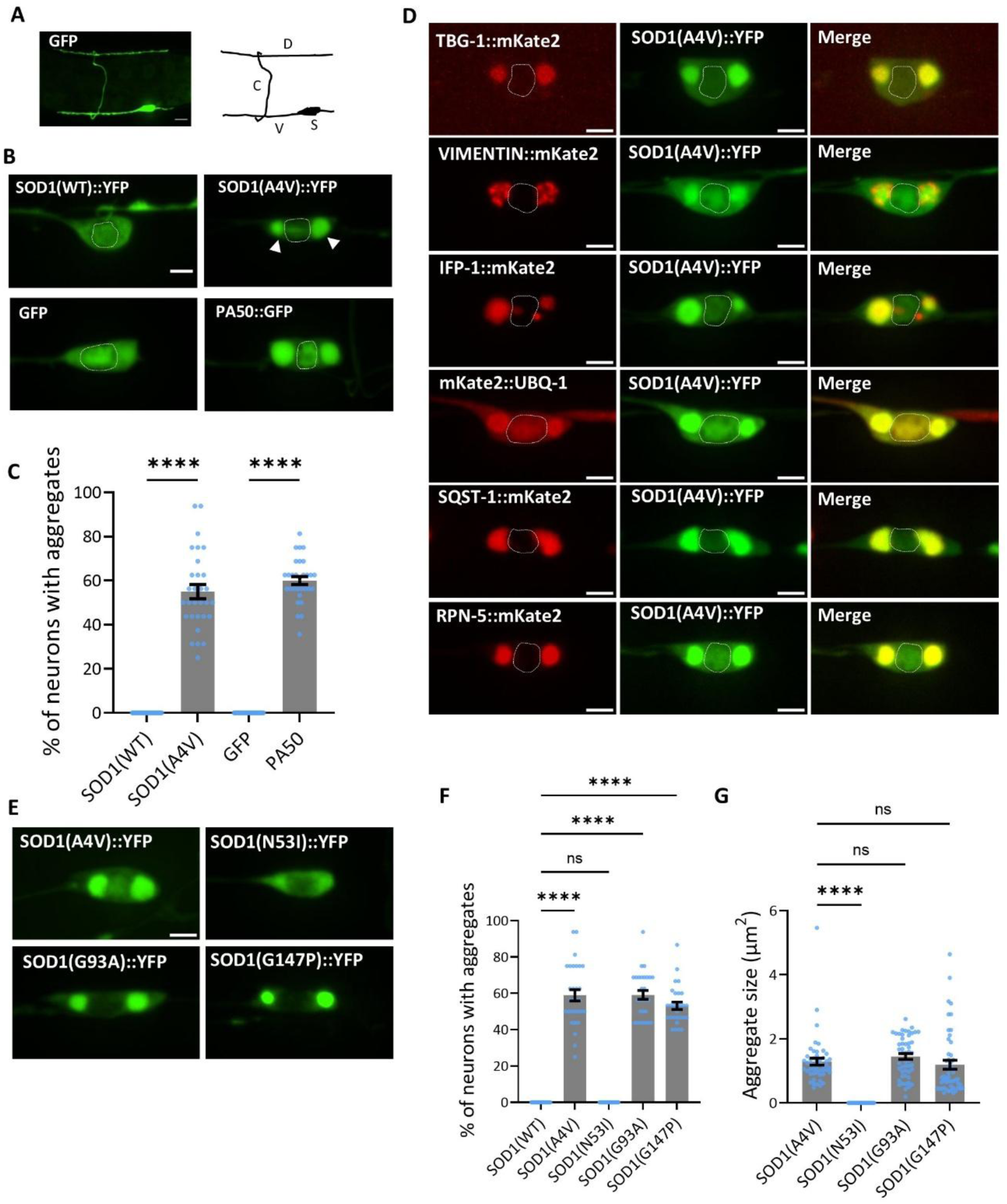
Neurodegenerative disease-associated misfolded proteins are sequestered into aggresomes in young *C. elegans* neurons. **A.** Morphology of a D-type GABAergic motor neuron labelled with GFP. These neurons have their soma (S) situated on the ventral side and a neurite that extends first ventrally (V), then circumferentially in a commissure (C), and finally dorsally (D). Scale bar = 4 µm. **B.** SOD1(A4V) and PA50 form aggregates in GABAergic motor neurons. Dashed lines outline the nuclei. White arrows indicate perinuclear aggregates. Scale bar = 2 µm. **C.** Percentage of neurons displaying aggregates upon expression of disease-associated SOD1 and PA50. N = 30, ****p < 0.0001, one-way ANOVA with Tukey’s post hoc test. **D.** SOD1(A4V) aggregates in GABAergic motor neurons colocalize with aggresome markers gamma tubulin (*C. elegans* TBG-1), intermediate filament (Mammalian Vimentin and *C. elegans* IFP-1), ubiquitin (*C. elegans* UBQ-1), p62 (*C. elegans* SQST-1), and proteasome subunit (*C. elegans* RPN-5). Dashed lines outline the nuclei. Scale bar = 2 µm. **E.** Aggresome formation correlates with SOD1 oligomeric state and cytotoxicity. Mutations A4V, G93A, and G147V, which form more trimers and exhibit higher cytotoxicity, induce aggresome formation. Mutation N53I, with fewer trimers and lower cytotoxicity, does not induce aggresomes. Scale bar = 2 µm. **F.** Probability of aggresome formation in different SOD1 mutations. N ≥ 28, ****P < 0.0001, one-way ANOVA with Dunnett’s post hoc test. **G.** Quantification of aggresome size across different SOD1 mutations. N = 50, ****P < 0.0001, one-way ANOVA with Tukey’s post hoc test. Data are represented as mean ± SEM in **C**, **F**, **G**.

Misfolded proteins are typically ubiquitinated and sequestered into aggresomes, protein quality-control compartments that form at the microtubule-organizing centre (MTOC)^5,6^. To determine the nature of SOD1(A4V) and PA50-triggered aggregates, we examined several aggresome markers: microtubule-organizing center (MTOC) protein γ-tubulin (*C. elegans* TBG-1), intermediate filament (mammalian Vimentin and *C. elegans* IFP-1), ubiquitin (*C. elegans* UBQ-1), P62 (*C. elegans* SQST-1), and a proteasome subunit RPN-5. SOD1(A4V) aggregates colocalized with all these aggresome markers (Fig. 1D). PA50 also colocalized with SOD1(A4V) and with the MTOC marker TBG-1 (Fig. S1C). These results suggest that SOD1(A4V) and PA50 aggregates in young animals are aggresomes. In contrast, the dipeptide repeats GA50 formed solid fibrillar-like aggregates within GABAergic motor neurons (Fig. S1D-G) that did not colocalize with the MTOC marker TBG-1 (Fig. S1C). Taken together, these results suggest that the misfolded proteins SOD1(A4V) and polyPA are sequestered into aggresomes in young *C. elegans* neurons.

To investigate if aggresome formation is associated with protein toxicity, we generated strains carrying SOD1 mutations with different oligomeric states and cytotoxicity. We found that more toxic mutations, including the trimer-stabilizing mutant G147P and ALS-associated mutants A4V and G93A^13,14^, triggered the formation of aggresomes (Fig. 1E-G), whereas the less toxic mutant N53I^13,14^ did not. These results suggest that neurons preferentially sequester more cytotoxic SOD1 proteins into aggresomes.

### Aggresomes are gradually damaged under sustained proteotoxic stress during aging

To examine how aggresome function changes with aging, we imaged aggresomes at different stages, from the L1 larval stage to old adult at day 13 (D13). We found that aggresomes were intact in younger animals (L1-L4) but gradually lost their integrity during aging (Fig. 2A, middle and lower panels, 2B). By D3 stage, aggresomes became indistinct shapes with blurry cores (Fig. 2A, middle and lower panels), and misfolded proteins were no longer accumulated within these structures. These observations indicate that the aggresomes become dysfunctional under persistent proteotoxic stress during aging. PA50-associated aggresomes showed similar deficits with aging (Fig. S2A). These impaired aggresomes were structurally different from autophagosomes or lysosomes, as they did not fully colocalize with the autophagosome marker LGG-1 or the lysosome marker LMP-1 (Fig. S2B), and the residual MTOC structure, marked by TBG-1, was still observed at this stage (Fig. 2B).

**Fig. 2:**
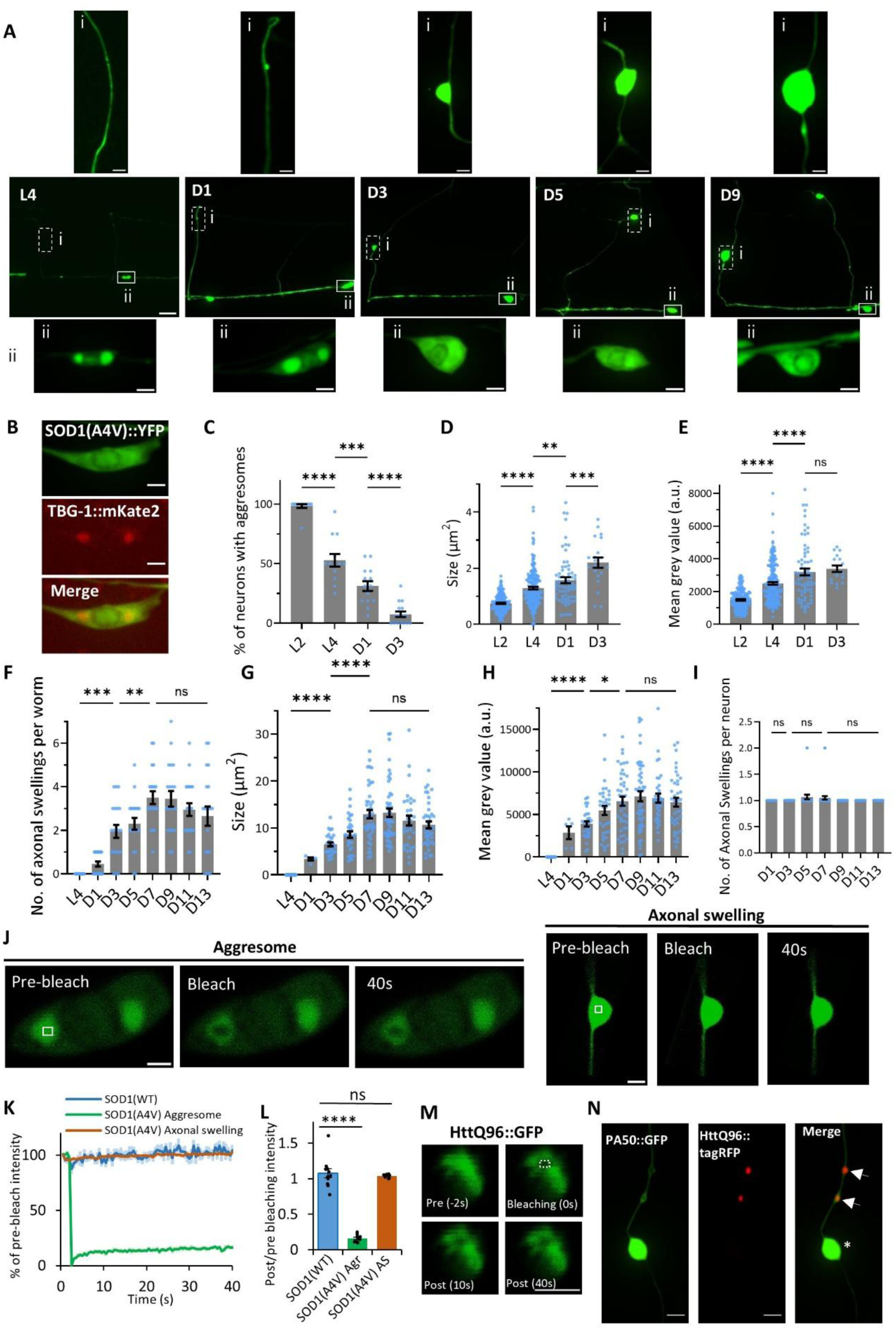
Misfolded proteins are sequestered into different compartments during aging. **A.** SOD1(A4V)::YFP is sequestered into aggresomes and axonal swellings during aging. Aggresomes in neuronal somas gradually lose integrity while axonal swellings at commissures form during aging. Dashed box (i) indicates axonal swelling (upper panel), solid line box (ii) indicates soma (lower panel); Scale Bars: 2 μm (upper and lower panels), 5 μm (middle panels). **B.** Colocalization of impaired aggresomes with TBG-1 at D3. Scale bars = 2 μm. **C.** Quantification of neurons with aggresomes during aging. Statistical comparisons: L1 vs. L4 (P<0.0001), D1 vs. D3 (P<0.0001), L4 vs. D1 (P=0.0005), n ≥ 13, ***P<0.001, ****P < 0.0001 from one-way ANOVA with Tukey’s post hoc test. **D**-**E**. Quantification of aggresome size (**D**) and fluorescent intensity (**E**) during aging. Statistical comparisons: L2 vs. L4 (P<0.0001), L4 vs. D1 (P=0.0041), D1 vs. D3 (P=0.0002), n ≥ 21, **P < 0.01, ****P < 0.0001, one-way ANOVA with Tukey’s post hoc test. **F**. Quantification of axonal swellings per animal during aging. Statistical comparisons: L4 vs. D3 (P=0.0001), D3 vs. D7 (P=0.0054), D7 vs. D13 (P=0.4371), n = 20, **P < 0.01, ****P < 0.0001, one-way ANOVA with Tukey’s post hoc test. **G**-**H**. Quantification of axonal swelling size (**G**) and intensity (**H**) during aging. Statistical comparisons: D3 vs. D7 (P=0.0102), D7 vs. D13 (P=0.3881 in **G**, >0.9999 in **H**), other comparisons P<0.0001, n = 20 worms, *P < 0.05, ****P < 0.0001, one-way ANOVA with Tukey’s post hoc test. **I.** Quantification of axonal swellings per neuron during aging. Statistical P > 0.9999, 0.7105, 0.6177 from left to right, n = 20 worms, one-way ANOVA with Tukey’s post hoc test. **J.** FRAP analysis of SOD1(A4V)::YFP mobility in aggresomes and axonal swellings. Left panels show aggresomes before and after photobleaching, and right panels show axonal swellings. Fluorescence in axonal swellings recovers rapidly, exceeding the imaging speed (2-3 Hz). Scale bar = 2 μm. **K.** Fluorescence changes before and after photobleaching in SOD1(WT) soma, SOD1(A4V) aggresomes, and axonal swellings. N ≥ 10. **L.** Quantification of fluorescence recovery percentage (40 s after bleaching). Statistical comparisons: SOD1(WT) soma vs. SOD1(A4V) aggresome (P < 0.0001), SOD1(WT) soma vs. SOD1(A4V) axonal swelling (P = 0.7994), n ≥ 10, ****P < 0.0001, one-way ANOVA with Tukey’s post hoc test. **M.** Photobleaching of HttQ96::GFP. Images were shown at −2 s, 0, 10 s and 40 s. Photobleaching was performed at 0 s within the region indicated by the dashed box. HttQ96::GFP exhibited solid-like properties. Scale bar = 2 µm. **N.** Location of PA50::GFP and httQ96::tagRFP. Note that PA50::GFP accumulated in the torpedo-like axonal swelling (white asterisk), not colocalized with the small beading-like swellings (white arrows) filled with httQ96 solid aggregates. Scale bar = 2 μm. Data are represented as mean ± SEM in **C-I, K-L**.

While the percentage of neurons with intact aggresomes decreased with age (Fig. 2C), the size and brightness of aggresomes that remained intact increased from L1 to D3 (Fig. 2D-E; Fig. S3A), suggesting continuing sequestration of misfolded proteins before aggresome dysfunction.

### A soluble misfolded proteins-induced axonal swelling sequesters pathogenic proteins following aggresome dysfunction

As aggresomes became dysfunctional, misfolded proteins were no longer accumulated in soma compartments. Instead, they were sequestered into an elliptic, torpedo-like swelling structure in neurites, from D3 through D13 (Fig. 2A middle and top panels, 2F). These structures were about 4-6 µm in length and 2-4 µm in width. They were much larger than aggresomes (Fig. 2D, G, Fig. S3A-B) and contained a significantly higher concentration of SOD1(A4V) (Fig. 2E, H, Fig. S3A-B). Axonal swellings were located in the axonal commissure, with typically one present per neuron (Fig. 2I), and were clearly distinguishable from the beading-like swellings^15^ associated with neuronal death (Fig. S3C). Other misfolded proteins, such as PA50, were also sequestered into similar structures after aggresome impairment (Fig. S3D).

Notably, the emergence of axonal swellings was caused by persisting proteotoxic stress during aging. It was not caused by a change in promoter activity during aging, as SOD1(WT) levels remained unchanged at D3 (Fig. S3E). Instead, it resulted from sustained proteotoxic stress and the progressive accumulation of misfolded SOD1(A4V) proteins within neurons during aging (Fig. S3E), likely due to age-related declines in other quality-control pathways, such as autophagy and proteasomal degradation. Without the progressive proteotoxic stress from the misfolded protein SOD1(A4V), the normal form of SOD1(WT) only induced very low-level age-dependent axonal swellings (Fig. S3F). Conversely, without aging, even high levels of misfolded proteins that impaired aggresome function at the L4 stage (Fig. S3G) were insufficient to induce axonal swelling formation at this stage (Fig. S3H).

Axonal swellings went through two stages: the initial growth stage (D1-D7), where both size and brightness increased, followed by a maintenance stage (D7-D13), where they stabilized or slightly decreased (Fig. 2F-H, Fig. S3B). The length/width ratio of the swellings was relatively stable from D1 to D13 (Fig. S3I-L), indicating well-maintained structures during aging.

Axonal swelling formation was accompanied by aggresome dysfunction. To examine if misfolded proteins from the dysfunctional aggresomes were transported to axonal swellings, we performed a photoactivation experiment by fusing SOD1(A4V) with UV-activated PA-GFP. We used RPN-5::mk2 as an aggresome marker, and selectively activated SOD1(A4V)::PA-GFP in the aggresome region with 405 nm light at the L4 stage (Fig. S4A). Prior to activation, SOD1(A4V)::PA-GFP fluorescence was dim in both the soma and neurites (axonal commissures). After activation, green fluorescence in aggresomes became bright (Fig. S4B-C). Four days later (D4), as aggresomes became dysfunctional, green fluorescence in the soma decreased (Fig. S4B, C), whereas the signal in axonal swellings significantly increased (Fig. S4B, D). This indicates that misfolded proteins were redirected from aggresomes (during impairment) to axonal swellings.

We then conducted FRAP (Fluorescence Recovery After Photobleaching) experiments to assess the mobility of misfolded proteins. SOD1(A4V) in aggresomes at L4 stage displayed low mobility with ∼20% recovery in 40 seconds (Fig. 2J-L), which aligns with previous reports in mammalian systems^16–18^. In contrast, SOD1(A4V) in axonal swellings at the D3 stage was highly mobile, with no difference from the non-aggregated SOD1(WT) (Fig. 2J-L). These results suggest that SOD1(A4V) proteins are in different physical states depending on the compartment in which they are sequestered. In young animals, misfolded SOD1 proteins are sequestered into aggresomes, where they form aggregates with low mobility. However, as the animals age and aggresomes become impaired, these misfolded proteins disperse and are redirected to axonal swellings in a soluble form. We named these elliptic, torpedo-like axonal swellings “SolAS” (<u>Sol</u>uble Misfolded Proteins-induced <u>A</u>xonal <u>S</u>wellings) to distinguish them from other types of axonal swellings, such as those formed by polyQ-associated solid aggregates (HttQ96), that accumulated in beading-like small swellings rather than in SolAS in the same neurite (Fig. 2M-N).

### Misfolded proteins are hierarchically sequestered into aggresomes and axonal swellings (SolAS) during aging

To understand how misfolded proteins are sequestered into different compartments during aging, we tracked misfolded proteins within the same neuron at different ages and used a Hidden Markov Model (HMM) to analyse compartment transitions. HMM is a statistical model that represents systems with hidden states, where observable outputs depend probabilistically on these states^19,20^. Based on our observations, we considered four distinct hidden states for neurons with misfolded proteins: 1) Diffuse state (D), where SOD1(A4V)::YFP is distributed throughout the neuron with no visible aggresomes or SolAS; 2) Aggresome state (Agr), where aggresomes are present but no SolAS is visible; 3) Impaired aggresome state (IA), where aggresomes lose integrity but no SolAS is visible; and 4) Axonal swelling state (AS), where aggresomes are impaired and SolAS appear (Fig. 3A). Images were captured at key stages every 36-48 hours: L1, L4, D1, D3 and up to D11 (Fig. 3B). After assigning each neuron to one of the four states at each stage, we calculated the state transition probabilities and mapped the state transition routes during aging.

**Fig. 3:**
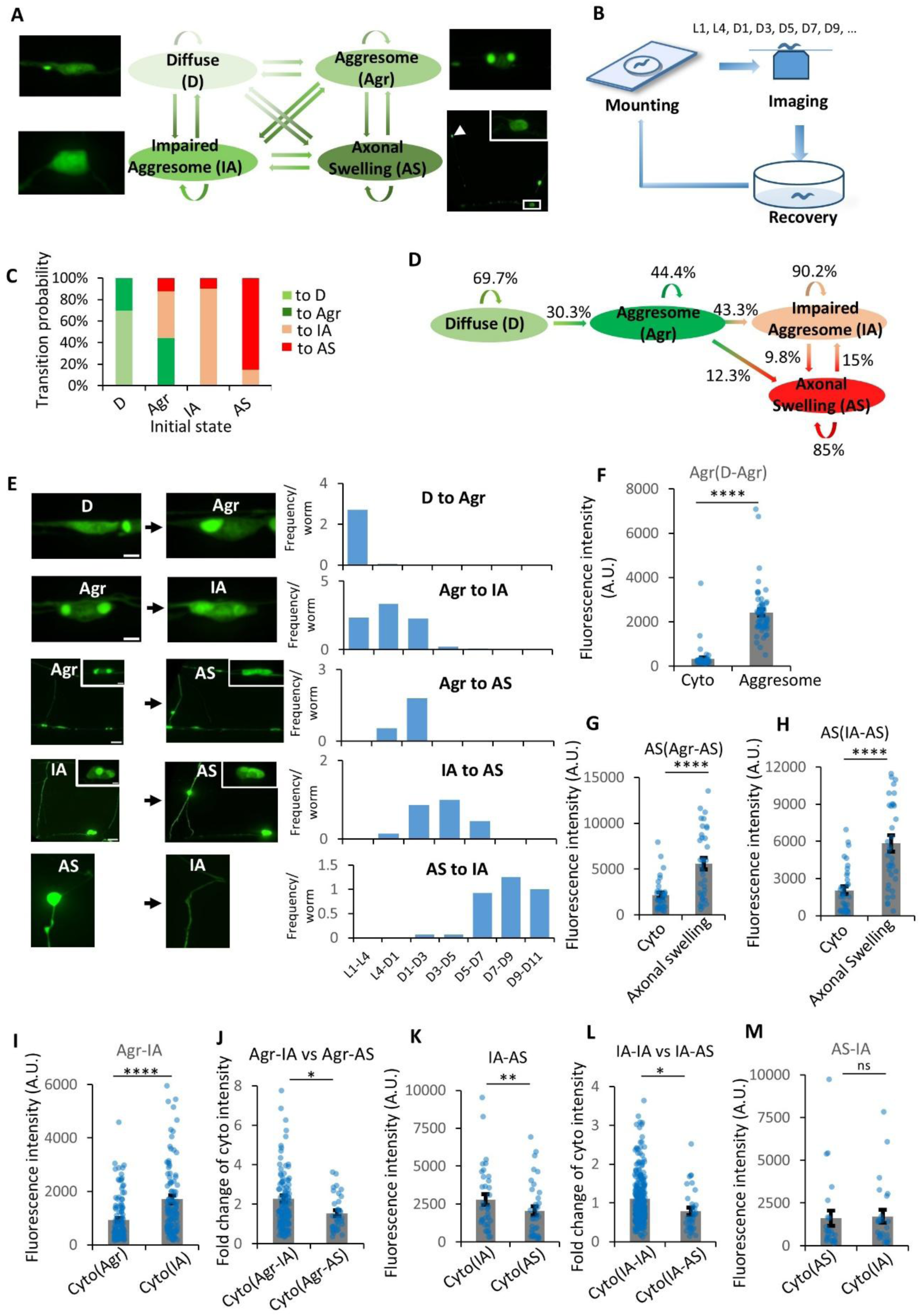
Hierarchical sequestration of misfolded proteins into aggresomes and axonal swellings during aging. **A.** HMM for the transitions between different neuron states in aging *C. elegans*. **B.** Schematic of longitudinal imaging of the same neurons in *C. elegans* over time. **C**-**D**. Transition probabilities (**C**) and transition routes (**D**) derived from the HMM model. Transitions between the four states (D, Agr, IA, AS) were quantified and transition probabilities were calculated by pooling data across different stages of aging. N = 305 (D), 282 (Agr), 511 (IA), 233 (AS). **E**. Aging-related state transitions in neurons with SOD1(A4V). The left panels show example transitions of same individual neurons at different stages, while the right panels display the distribution of different transitions at different ages. N =58 (D-Agr), 123 (Agr-IA), 35 (Agr-AS), 36 (IA-AS), 26 (AS-IA). **F**-**G**. Fluorescence intensity of SOD1(A4V)::YFP in the same neurons during the transition from diffuse to aggresome (D-Agr) (**F**) and from aggresome to axonal swelling (Agr-AS) (**G**). ****P < 0.0001 from paired t test, n = 58 (**F**), 35 (**G**). **H.** Fluorescence intensity changes within the same neurons during the transition from impaired aggresome to axonal swelling (IA-AS). ****P < 0.0001 from paired t test, n = 36. **I.** Cytoplasmic fluorescence intensity in the same neurons following the transition from aggresome to impaired aggresome (Agr-IA). N = 123, ****P < 0.0001 from paired t test. **J.** Fluorescence intensity changes in the cytoplasm during transitions from aggresome to axonal swelling (Agr-AS) and from aggresome to impaired aggresome (Agr-IA). *P = 0.0301 from unpaired t test, n =123 (Agr-IA), n = 35 (Agr-AS). **K.** Cytoplasmic fluorescence intensity in the same neurons following the transition from impaired aggresome to axonal swelling (IA-AS). **P = 0.0012 from paired t test, n = 36. **L.** Fluorescence intensity changes in the cytoplasm during the transition from impaired aggresome to axonal swelling (IA-AS) and during the self-transition (IA-IA). *P = 0.0223 from unpaired t test, n = 327 (IA-IA), n = 36 (IA-AS). **M.** Fluorescence intensity changes during the reverse transition from axonal swelling to impaired aggresome (AS-IA). P = 0.6241 from paired t-test, n = 26. Data are represented as mean ± SEM in **F**-**M**.

The state transition routes revealed that misfolded proteins were hierarchically sequestered into different compartments (Fig. 3C, D). Specifically, neurons in the diffuse state (D) transitioned to the aggresome state (Agr) with a probability of 30.3%. From the aggresome state (Agr), neurons either transitioned directly to the axonal swelling state (AS) with a probability of 12.3% (fast transition) or transitioned first to the impaired aggresome state (IA) with a probability of 43.3%, before subsequently transitioning to the axonal swelling state (AS) with a probability of 9.8% (slow transition) (Fig. 3C, D). The delay observed in some neurons remaining in the IA before transitioning to the AS may reflect variability in the accumulation of sufficient misfolded proteins needed to trigger axonal swelling formation. Nevertheless, they both underwent similar transitions from intact aggresomes to impaired aggresomes in the soma. The AS state could revert to the IA state with a probability of 15% (Fig. 3C, D). These results confirm that misfolded proteins are initially sequestered into somatic aggresomes, and as aggresomes deteriorate with age, SolAS emerge to further sequester misfolded proteins.

To investigate how this hierarchical pathway evolves with animal age, we tracked state transitions in animals of different ages. Our results showed a transition from the diffuse to the aggresome state (D-Agr) during early larval stages from L1 to L4 (Fig. 3E, Fig. S5A). Then, a transition from the aggresome to the impaired aggresome state (Agr-IA) occurred, spanning from L1 to D3 and peaking at L4-D1. The transition from the aggresome state to the axonal swelling state (Agr-AS) reached its peak at D1-D3, and the transition from impaired aggresome state to the axonal swelling state (IA-AS) peaked later at D3-D5. Interestingly, in mid-to-late age (D7 to D11), axonal swellings could revert to the impaired aggresome state (AS-IA) (Fig. 3E, Fig. S5B-D). Except for the state transition, the maintenance of self-state, such as aggresomes (Agr-Agr), was associated with increasing size and intensity, suggesting the persistence of proteostatic stress (Fig. S5E, F). These results suggest a dynamic sequestration mechanism that shifts from aggresomes to SolAS under proteotoxic stress with age.

### SolAS function as quality-control compartments that reduce somatic misfolded protein burden

We next sought to understand the functional differences between aggresomes and SolAS. By quantifying SOD1(A4V)::YFP intensity in the soma cytoplasm, aggresomes, and neurite SolAS across different states and transitions, we found that neurons transitioning from the diffuse to aggresome state (D-Agr), the SOD1(A4V)::YFP intensity in aggresomes was significantly higher than in the cytoplasm (Fig. 3F), confirming aggresomes as quality-control compartments to sequester toxic proteins and reduce cytoplasmic misfolded protein levels. Similarly, in neurons transitioning to SolAS (Agr-AS or IA-AS), the SOD1(A4V)::YFP intensity in SolAS was also significantly higher than in the cytoplasm (Fig. 3G, H), indicating that SolAS serve as quality-control compartments to sequester intracellular misfolded proteins after aggresome dysfunction.

Compared to aggresomes (∼2 μm², Fig. 2D), the size of SolAS was significantly larger (10–14 μm², Fig. 2G). Consistent with this, the fluorescence intensity of SOD1(A4V)::YFP in SolAS was much higher than in aggresomes (Fig. 2E, H; Fig. 3F-H), indicating that SolAS can accommodate a greater burden of misfolded proteins. The higher cytoplasmic levels of toxic proteins in neurons with SolAS, compared to those with aggresomes, suggest that SolAS are high-threshold compartments for sequestration (Fig. 3F-H).

To understand how SolAS reduce soma intracellular misfolded protein levels, we measured the cytoplasmic SOD1(A4V)::YFP intensity in neurons with and without SolAS formation. Impaired aggresomes led to dispersed misfolded proteins in the soma, as shown by nearly a two-fold increase in cytoplasmic SOD1(A4V) levels in neurons shifting from the functional aggresome state to the impaired aggresome state (Agr-IA) (Fig. 3I). However, as axonal swellings formed, either through the fast transition route (Agr-AS) or the slow route (IA-AS), levels of cytoplasmic SOD1(A4V) dropped significantly (Fig. 3J–L), confirming that SolAS as quality control compartments to reduce soma misfolded protein levels. Interestingly, some of the SolAS might be removed from neurites without an increase in intracellular SOD1(A4V) level (AS-IA) (Fig. 3M). These findings indicate that SolAS formation restricts the buildup of dispersed SOD1(A4V) proteins in the soma.

### SolAS are precisely positioned and well-organized structures in neurites

SolAS are located at axonal commissures, which extend circumferentially from the ventral to the dorsal, between the body wall muscles and the large multinucleated hypodermis (skin)^21,22^. The hypodermis forms a thin sheet across the muscles and expands into large lateral ridges between dorsal and ventral muscles. By labelling these surrounding cells with different fluorescent markers, we found that SolAS were confined to the segment crossing the lateral hypodermal ridge, embedded within this tissue, and was never observed in the hypodermal sheet region (Fig. 4A, B). The occurrence of SolAS in the lateral ridge region decreased with increasing distance from the ventral side (Fig. 4B, C), suggesting a precise positioning of SolAS within the commissure.

**Fig. 4:**
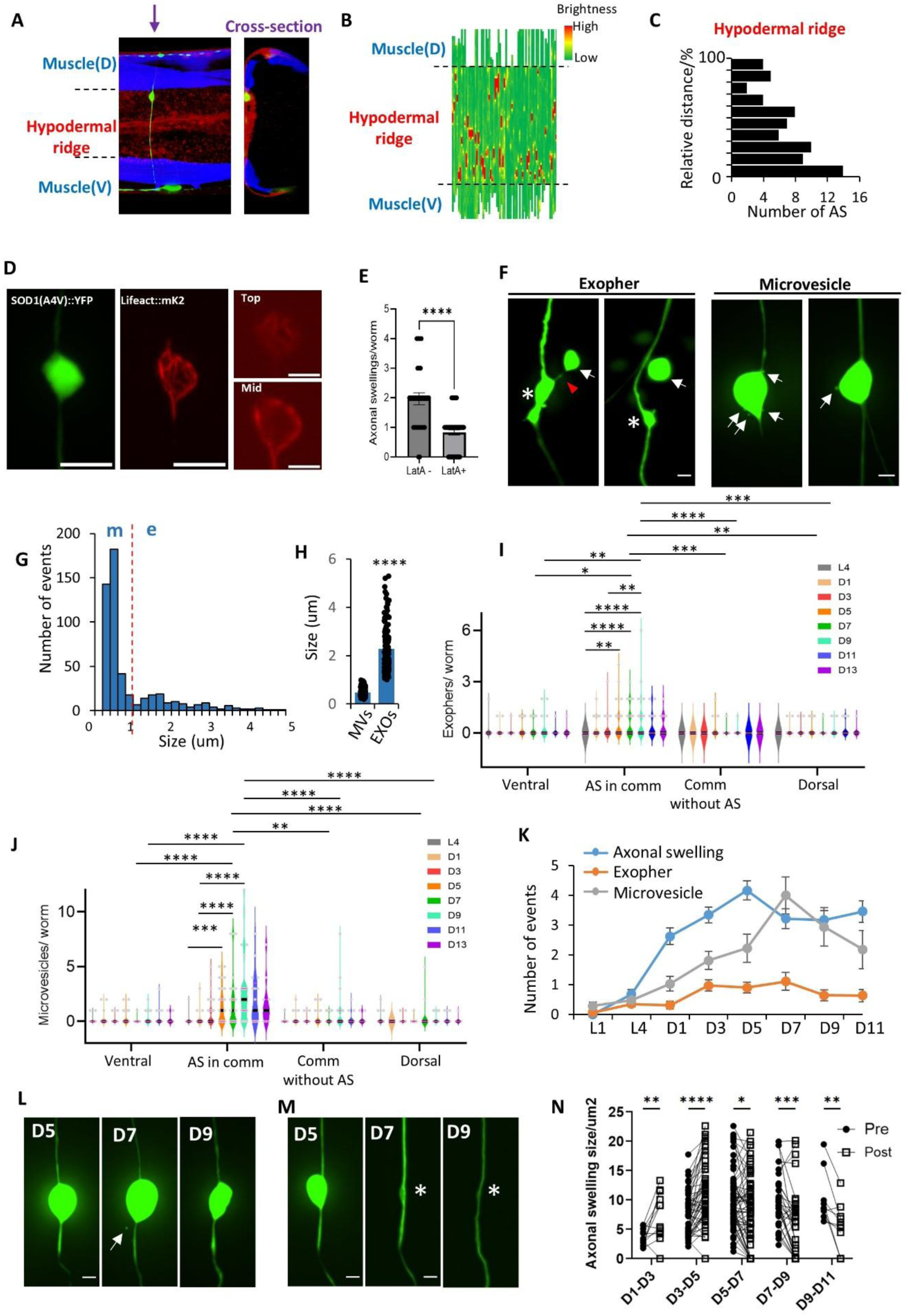
Axonal swellings are precisely positioned and organized structures to expel soluble toxic proteins via microvesicles and exophers. **A.** Location of axonal swellings. Arrow indicates the cross-section site. Scale bar = 2 µm. **B**-**C**. Heatmap (**B**) and histogram (**C**) of axonal wellings distribution along axonal commissures. Red signals (high fluorescence intensity) mark axonal swellings (**B**). The size of the commissure segment spanning the hypodermal ridge is normalized to 1. The reference point 0 in (**C**) is set at the lower dashed line in (**B**). N=69. **D.** A cage-like actin network as the structural scaffold for SolAS. Actin network was labelled with lifeact::mk2. Scale bar = 4 µm (left two panels), 2 µm (right two panels). **E.** Quantification of axonal swellings in worms treated with actin polymerization inhibitor Latrunculin A (LatA). N = 30 (LatA-), 65 (LatA+), ****p < 0.0001 from unpaired t test. **F.** Exophers and microvesicles expelled from axonal swellings. Left panel: white asterisks indicate axonal swellings after exopher expulsion; white arrows highlight exophers; red arrow points to the thin tube commonly associated with exopher generation. Right panel: white arrows mark microvesicles near or buding from axonal swellings. Scale bar = 2 µm. **G**-**H**. Histogram (**G**) and quantification (**H**) of the size of vesicles expelled from axonal swellings. Two distributions were observed, with a size cutoff of 1.0 µm to differentiate microvesicles (m) from exophers (e). Bin size = 0.2 µm, n = 517, ****P < 0.0001 from unpaired t-test. **I**-**J**. Violin plot of exophers (**I**) and microvesicles (**J**) distributions at different parts of a neuron during aging. N = 31 (L4), 31 (D1), 32 (D3), 38 (D5), 31 (D7), 28 (D9), 17 (D11), 11 (D13). **I.** Statistical comparisons within “AS in comm” group: L4 vs. D5 (P=0.0045), L4 vs D7 (P<0.0001), L4 vs. D9 (P<0.0001), D3 vs. D9 (P=0.0016); Statistical comparisons within D7 group: AS in comm vs. Ventral (P=0.0481), AS in comm vs. Comm without AS (P=0.0001), AS in comm vs. Dorsal (P=0.0001); Statistical comparisons within D9 group: AS in comm vs. Ventral (P=0.0078), AS in comm vs. Comm without AS (P<0.0001), AS in comm vs. Dorsal (P=0.0002); **J.** Statistical comparisons within “AS in comm” group: L4 vs. D5 (P=0.0002), D1 vs D7 (P<0.0001), D1 vs. D9 (P<0.0001); Statistical comparisons within D7 group: AS in comm vs. Ventral (P<0.0001), AS in comm vs. Comm without AS (P<0.0001), AS in comm vs. Dorsal (P<0.0001); Statistical comparisons within D9 group: AS in comm vs. Ventral (P<0.0001), AS in comm vs. Comm without AS (P<0.0001), AS in comm vs. Dorsal (P<0.0001); *P < 0.05, **P < 0.01, ***P < 0.001, ****P < 0.0001 from one-way ANOVA with Tukey’s post hoc test. Thick dark lines represent the median, magenta lines denote the quartiles, and data points are labeled in grey. **K.** Quantification of axonal swellings, exophers, and microvesicles during aging. N number same as in **I**-**J**. **L.** Shrinking axonal swellings during aging were accompanied by the formation of microvesicles. White arrows point to microvesicles. Same axonal swelling imaged at D5, D7, and D9. Scale bar = 2 µm. **M.** Complete expulsion of an axonal swelling by vesicles. Asterisk indicates the site of extrusion. The neurite maintains a normal morphogy without any signs of degeneration after several days of expulsion. Scale bar = 2 µm. **N.** Quantification of axonal swelling size change at different time transitions (D1-D3, D3-D5, D5-D7, D7-D9, D9-D11). Dots and squares represent measurements from pre (or earlier) (e.g., D1) and post (or later) (e.g., D3) time points, respectively. Axonal swellings that are completely expelled by the later time are recorded with a size of 0. N = 12, 52, 63, 35, 10. P = 0.0092, 0.0001, 0.0236, 0.0002, 0.0066 from paired t test. Data are represented as mean ± SEM in **E**, **H**-**K**, **N**.

Maintaining the regular shape of the SolAS, which are filled with soluble misfolded proteins in the hypodermal ridge segment, requires supporting skeletons. We examined whether actin provides this support by labelling actin filaments with lifeact-mKate2. Our observations showed that the SolAS were surrounded by a cage-like actin network (Fig. 4D). We further found that this actin network was essential for SolAS formation, as inhibiting actin polymerization using latrunculin A (LatA) significantly decreased SolAS numbers (Fig. 4E). These results indicate that SolAS are well-organized structures in neurites.

### SolAS serve as hotspots for expelling misfolded proteins via exophers and microvesicles

Canonical quality-control compartments, such as aggresomes, not only sequester misfolded proteins but also serve as hubs for their clearance by autophagy. We then sought to test whether SolAS similarly mediate the clearance of misfolded proteins. The hypodermal cells in *C. elegans* have strong engulfing functions, similar to microglial cells in mammals^23–25^. Given that SolAS are embedded within the hypodermis, we hypothesized that they may extrude misfolded proteins that are subsequently engulfed by the hypodermis. We quantified SOD1(A4V)::YFP–positive vesicles in the vicinity of SolAS and observed vesicles of varying sizes that filled with SOD1(A4V)::YFP, either closely associated with or budding from the SolAS. The vesicle size distribution showed two populations: small vesicles around 500 nm and larger ones from 1 to 5 µm (Fig. 4F-H). Notably, the larger vesicles appeared to be linked by a thin tube (Fig. 4F), a typical feature of exophers. These structures mediate the extrusion of neural waste from the somata of touch receptor neurons in *C. elegans* and mammalian neuron^26,27^. Interestingly, we found that exophers can be generated from SolAS at neurites (Fig. 4F-H). The smaller vesicles, with a size range from 200 nm to 1 µm (average 500 nm), appeared to bud directly from SolAS (Fig. 4F-H), resembling microvesicles. The microvesicles are distinct from the starry signals of hypodermal vesicles in *C. elegans*^24^. First, their size and shape were more consistent (Fig. 4G), unlike the varied sizes and shapes of the starry vesicles. Second, the microvesicles were closely associated with SolAS, often appearing to bud from them (Fig. 4F), whereas starry vesicles were not directly connected to commissures or swellings. Interestingly, exopher vesicles were expelled along with the actin network from SolAS. In contrast, microvesicles did not have an actin sheath (Fig. S6A-B).

To further confirm the relationship between these vesicles and SolAS, we quantified their distribution along neurons at three regions: the ventral side (including the soma), the commissure with SolAS, and the dorsal side. Our findings indicated that exophers and microvesicles were notably more prevalent in the SolAS region than in other areas during aging (Fig. 4I, J), supporting the idea that SolAS serve as hotspots for exophers and microvesicles. In addition, exophers and microvesicles appeared after SolAS formation (Fig. 4K). When tracking individual SolAS over time, we found that vesicle expulsion often led to a reduction in SolAS size or its complete removal from the commissure (Fig. 4L-N). These vesicles were initially bright in green fluorescence and disappeared over time, likely due to degradation in hypodermal cells (Fig. 4L). Remarkably, even after expelling large amounts of toxic proteins, the commissures remained structurally intact, with no signs of degeneration (Fig. 4M), suggesting that SolAS are different from degeneration- or death-associated axonal swellings. This also supports our HMM model that axonal swelling states can revert to impaired aggresome states (AS-IA) after expelling toxic protein content (Fig. 3D, E), without increasing misfolded protein levels in the soma (Fig. 3M). Extrusion of exophers and microvesicles reduces SolAS size, starting at the D5-D7 transition (Fig. 4N). Collectively, these results suggest that SolAS function as a key site to expel misfolded proteins through exophers and microvesicles, thereby reducing intracellular misfolded protein levels.

### SolAS do not disrupt microtubules or axon transport

To assess whether SolAS disrupt neural function, we examined microtubule organization and axonal transport. Degeneration-associated axonal swellings impair axonal transport with microtubule looping^28^. However, microtubules labelled with microtubule-binding domain of ensconsin (EMTB) remained well-organized within SolAS at different ages (D2 and D5), and no microtubule looping was observed (Fig. S7A).

In addition, normal microtubule function is essential for SolAS formation, as disrupting the microtubule system with nocodazole significantly reduced the number of SolAS (Fig. S7B). This likely indicated the need for microtubule-based antegrade transport to redirect SOD1(A4V) from the soma to SolAS. Consistent with this idea, mutants of *unc-116* (kinesin-1 heavy chain KIF5 in mammals) showed defects in SolAS formation, and reintroducing UNC-116 specifically in these neurons rescued SolAS formation defects (Fig. S7C). These results suggest that SolAS formation depends on kinesin-mediated anterograde transport.

We also assessed SolAS impact on axonal transport by examining synaptic vesicle cargo with SNB-1::mKate2, a presynaptic protein that is transported anterogradely through the commissure to the dorsal side in DD neurons^29^ (Fig. S7D). We found no significant difference in the intensity and number of SNB-1 puncta at presynaptic regions at different age stages (D3 and D7) (Fig. S7E, F), indicating that SolAS do not significantly disrupt axonal transport. We also examined excitation propagation in DA and DB motor neurons using jRCaMP1b^30^. We found no significant difference in calcium signals between proximal and distal regions (Fig. S7G, H). All these findings indicate that SolAS do not significantly compromise axonal transport or excitation propagation, at least at the ages examined in this study.

### Ubiquitin tagging maintains aggresome integrity and suppresses the formation of SolAS during aging

The identification of the protective role of SolAS during aging prompted us to determine the molecular mechanisms underlying aggresome impairment and SolAS formation. Given the importance of ubiquitin for aggresome function and an increase in deubiquitination during *C. elegans* aging^31^, we studied the ubiquitin pattern in the same neuron at L4 and D3. We found that ubiquitin was colocalized with aggresomes at L4 and became dispersed before aggresome impairment during aging (Fig. 5A), suggesting that the loss of ubiquitination may precede and trigger SOD1 dispersion. To further test this idea, we fused ubiquitin to SOD1(A4V) so that ubiquitin could not be removed by deubiquitination. We found that this modification suppressed SOD1 dispersion and preserved aggresome integrity during aging, with aggresomes remaining intact even at D7(Fig. 5B-D). In contrast, SolAS formation was significantly reduced (Fig. 5E-F). This was further validated by fusing ubiquitin with PA50, which similarly maintained aggresome integrity and suppressed SolAS formation during aging (Fig. S8A-E). These results suggest that ubiquitin tagging of misfolded proteins is critical for preserving aggresome integrity and inhibiting SolAS formation during aging.

**Fig. 5:**
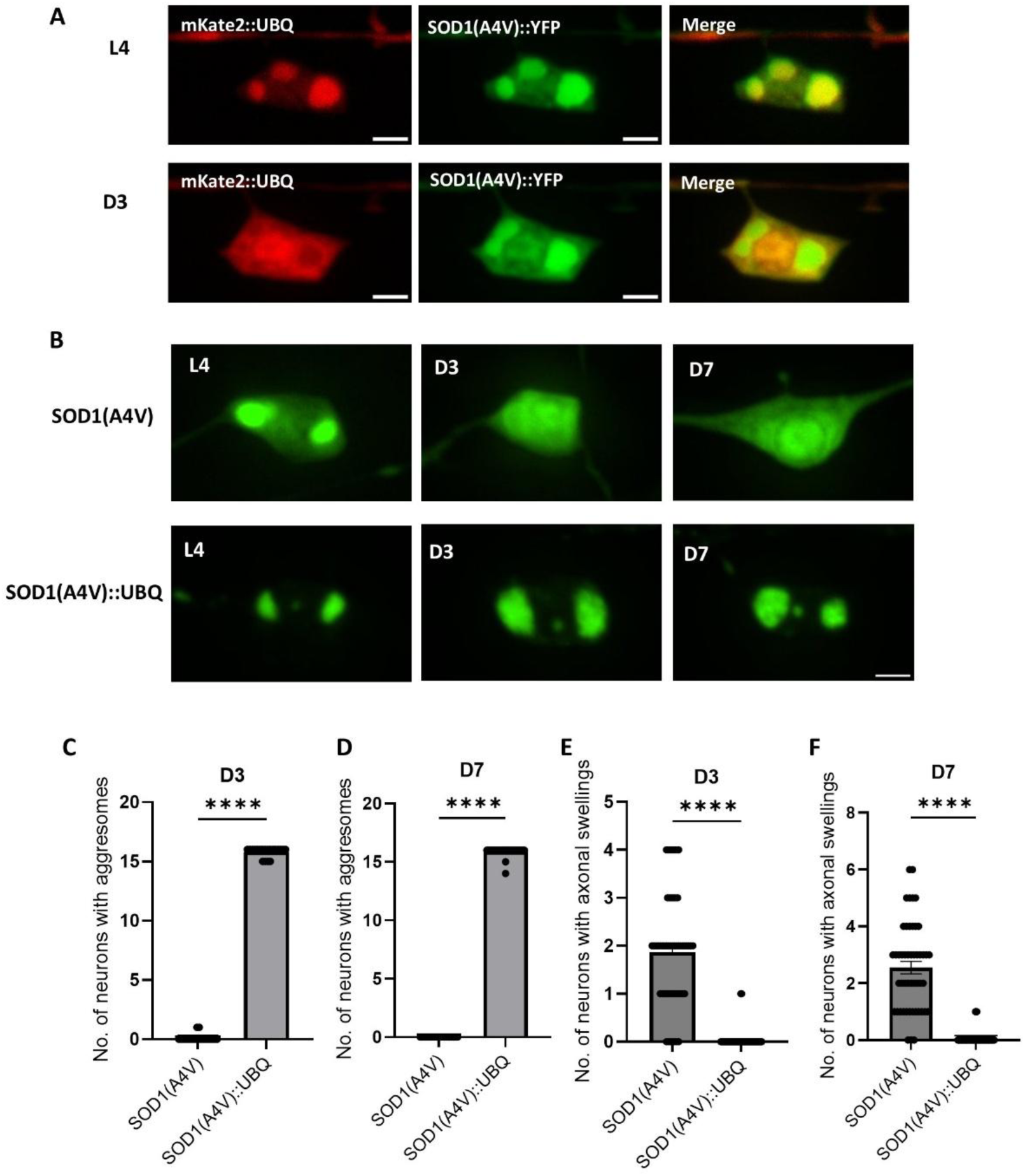
Ubiquitin signalling maintains aggresome integrity and suppresses SolAS formation during aging. **A.** Changes in the colocalization of ubiquitin with aggresomes before and after aggresome impairment. The upper panel shows mKate2::UBQ colocalized with aggresomes at L4, while the lower panel shows a hollow pattern of mKate2::UBQ in aggresomes and reduced colocalization at D3. The same neuron of the same worm was imaged at L4 and D3 stages. Scale bar = 2 µm. **B.** Fusion of ubiquitin with SOD1(A4V) mitigates aggresome impairment and preserves aggresome integrity during aging. Imaging was performed at L4, D3, and D7 stages. Scale bar = 2 µm. **C**-**D**. Quantification of neurons with aggresomes in animals with and without ubiquitin fusion to SOD1(A4V) at D3 (**C**) and D7 (**D**). N = 30 (**C**), 29 (**D**), ****P < 0.0001 from unpaired t-test. **E**-**F**. Quantification of neurons with axonal swellings in animals with and without ubiquitin fusion to SOD1(A4V) at D3 (**E**) and D7 (**F**). N = 54 (**E**), 30 (**F**), ****P < 0.0001 from unpaired t-test). Data are represented as mean ± SEM in **C**-**F**.

### SUMOylation of misfolded proteins promotes SolAS formation during aging

To identify molecules involved in SolAS formation, we screened a series of markers and mutants. We found that none of the aggresome markers tested, including TBG-1, IFP-1, SQST-1, and RPN-5, colocalized with SolAS (Fig. 6A), indicating that they are different from aggresomes at the molecular level. In our screen, we found that SUMO showed a strong signal in SolAS, compared with the notably lower ubiquitin level (Fig. 6A). Interestingly, aging has been associated with a decline of ubiquitination^32^ and an increase in SUMOylation^31^. To investigate this further, we fused SUMO (SMO-1 in C. elegans) to SOD1(A4V). This modification significantly decreased aggresome numbers in both young and old animals and enhanced SOD1(A4V) dispersion in the soma, even at the L4 stage (Fig. 6B, C). Importantly, SUMO fusion also increased SolAS formation at D3 (Fig. 6D, E), indicating that SUMOylation promotes SolAS formation while reducing aggresome formation.

**Fig. 6:**
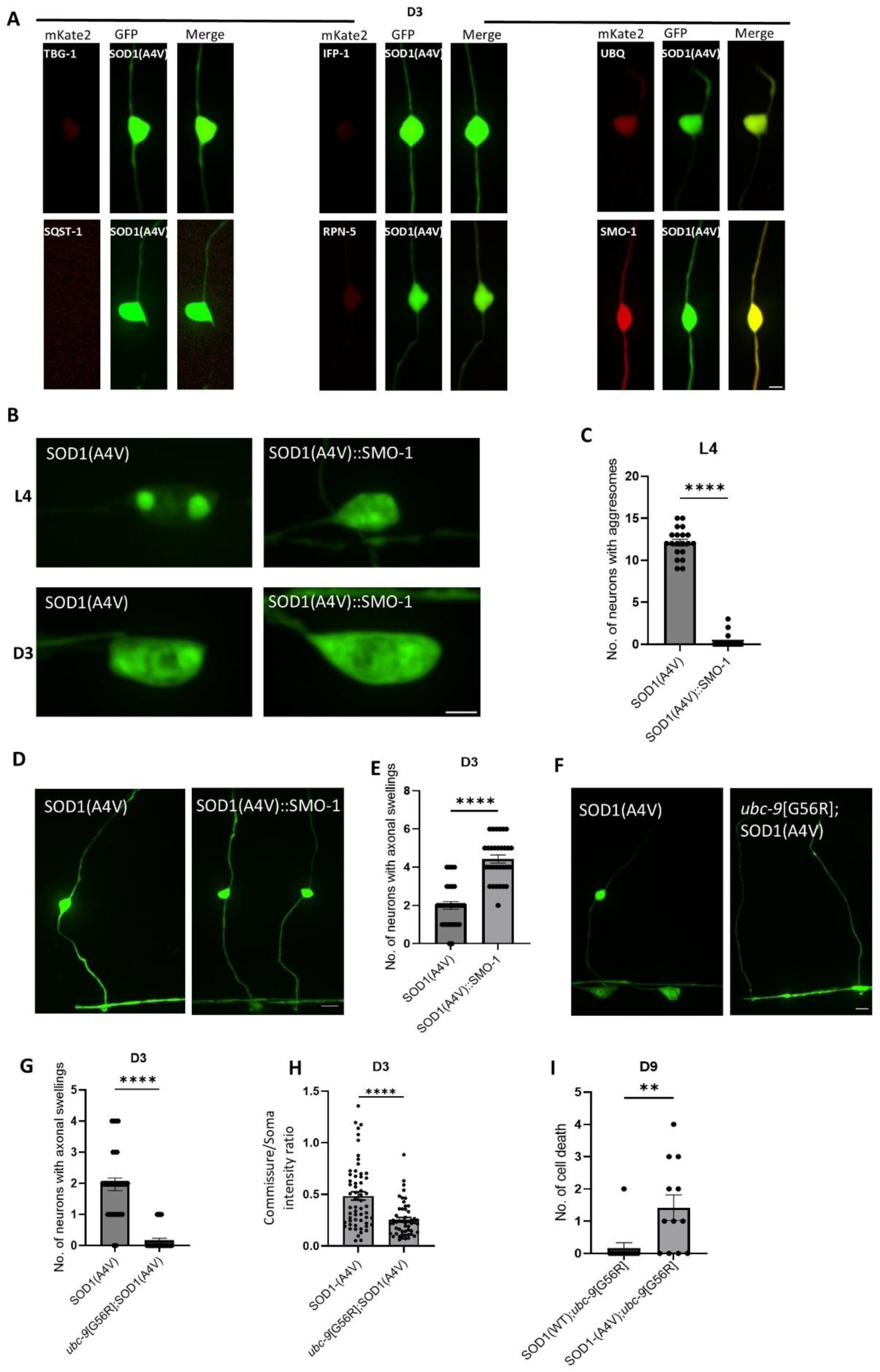
SUMO signalling promotes axonal swelling formation and suppresses aggresomes during aging. **A.** Aggresome markers are not colocalized with axonal swellings, but SMO-1 shows a much brighter signal within axonal swellings. Scale bar = 2 µm. **B.** Fusion of SMO-1 with SOD1(A4V) abolishes aggresome formation in young L4-stage animals. Scale bar = 2 µm. **C.** Quantification of aggresome numbers in L4-stage worms with or without SMO-1 fusion to SOD1(A4V). ****P < 0.0001 from unpaired t-test, n = 30. **D.** Fusion of SMO-1 with SOD1(A4V) promotes axonal swelling formation at D3. Scale bar = 5 µm. **E.** Quantification of axonal swelling numbers at D3 in worms with or without SMO-1 fusion to SOD1(A4V). N = 30, ****P < 0.0001 from unpaired t-test. **F.** Mutation of the E2 SUMO-conjugating enzyme UBC-9 suppresses the formation of axonal swellings. Scale bar = 5 µm. **G.** Quantification of axonal swelling numbers in *ubc-9*[G56R] temperature sensitive mutants. Experiments conducted at 23 ℃. N = 30, ****P < 0.0001 from unpaired t-test. **H.** Quantification of SOD1(A4V) distribution in the commissure and soma (intensity of commissure/soma) in *ubc-9* mutants. N = 52, ****P < 0.0001 from paired t-test. **I.** Quantification of neuron death in *ubc-9* mutants expressing either wild-type SOD1 or SOD1(A4V). Data from D9 animals. N = 12, **P =0.0084 from unpaired t-test. Data are represented as mean ± SEM in **C**, **E**, **G**-**I**.

SUMOylation involves three enzymes: E1 (SUMO-activating), E2 (SUMO-conjugating), and E3 (SUMO ligase). *C. elegans* has multiple E1 and E3 enzymes but only one E2 UBC-9^33^. We then generated a temperature-sensitive *ubc-9* mutant (G56R) using CRISPR gene editing^34^ and found that at restrictive temperatures (24 ℃) these mutants showed significantly reduced SolAS formation (Fig. 6F, G), confirming a positive role of SUMOylation in SolAS formation. In addition, *ubc-9* mutants displayed a lower commissure-to-soma fluorescence intensity ratio (Fig. 6H), indicating a defect in sequestering misfolded proteins to commissures. Notably, this resulted in increased neuron death in SOD1(A4V) animals (Fig. 6I), suggesting that disrupting the proper sequestration of misfolded proteins into SolAS leads to neurotoxicity and degeneration.

Next, we asked whether SUMO modification could induce SolAS formation with pathogenic proteins that typically do not form SolAS, such as polyGA, which forms solid amyloid plaques in mammals^35^. In *C. elegans*, GA50 formed similar solid fibrillar aggregates in neuron soma that persist during aging without forming SolAS (Fig. 7A-D, Fig. S1D, G). Interestingly, fusion of SMO-1 to GA50 suppressed amyloid GA50 aggregate formation in young animals (L4), converting them into soluble proteins (Fig. 7A, C), and promoted SolAS formation at D3 (Fig. 7B, D). Importantly, this conversion reduced GA50 toxicity, as the SMO-1 fusion significantly reduced neuronal death and extended animal lifespan (Fig. 7E, F). These findings indicate that SUMO fusion with solid amyloid proteins is sufficient to promote SolAS formation, facilitating their sequestration and reducing neurotoxicity.

**Fig. 7:**
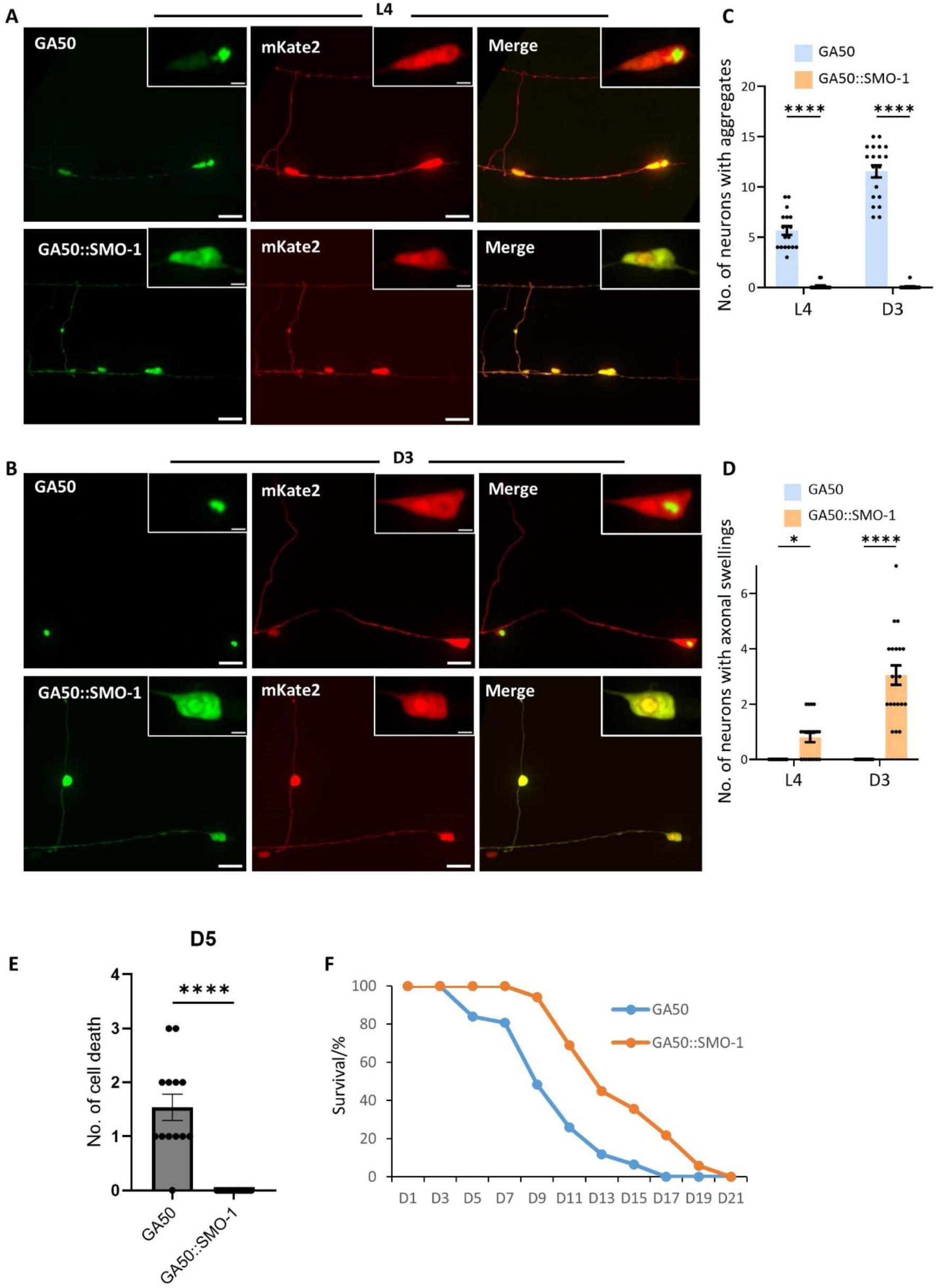
Conversion of toxic dipeptide repeat GA50 from solid amyloid aggregates in the soma to soluble aggregates in axonal swellings by fusion with SUMO. A-B. Images of GA50 with or without fusion to SMO-1 at the L4 (A) and D3 (B) stages, respectively. The green channel shows GFP fusion, and the red channel indicates neuron morphology using mKate2. Note that GA50 forms solid aggregates in the soma, while the fusion of SMO-1 with GA50 transforms these aggregates into a soluble state, driving the formation of axonal swellings by D3. Scale bar = 2 µm. **C**-**D**. Quantification of aggregates (**C**) and axonal swellings (**D**) in animals expressing GA50, with or without fusion to SMO-1, at both L4 and D3 stages. Data are presented as mean ± SEM, n ≥ 18, L4: GA50 vs. GA50::SMO-1 (P=0.0252), *P<0.05, ****P < 0.0001 from one-way ANOVA with Tukey’s post hoc test. **E**. Quantification of cell death in worms expressing GA50 or GA50::SMO-1 fusion at D5. Data are presented as mean ± SEM, n = 13, P < 0.0001 from unpaired t test. **F**. Survival curve of worms expressing GA50 or GA50::SMO-1 fusion. Mean survival time: 10.14 ± 0.34 (GA50), 14.43 ± 0.37 (GA50::SMO-1); Median survival time: 9 (GA50), 13 (GA50::SMO-1); n = 93 (GA50), 87 (GA50::SMO-1), p < 0.0001 from log-rank (Mantel-Cox) test.

### Ubiquitin-SUMO balance regulates the hierarchical sequestration of misfolded proteins into aggresomes and axonal swellings

To investigate how different posttranslational modifications regulate the sequestration of misfolded proteins into aggresomes and SolAS, we developed a technique to monitor ubiquitin-and SUMO-modifications on SOD1(A4V) based on split CFP-YFP complementation. The FP11 segment binds either CFP1-10 or YFP1-10, generating CFP or YFP fluorescence, respectively. Neither FP11, CFP1-10, or YFP1-10 alone produce fluorescence^36^. To detect ubiquitin- or SUMO-modifications on SOD1(A4V), we fused FP11 to SOD1(A4V), CFP1-10 to ubiquitin, and YFP1-10 to SUMO. If SOD1(A4V) is ubiquitinated, CFP fluorescence will be detected, whereas if it is SUMOylated, YFP fluorescence will be observed (Fig. 8A).

**Fig. 8:**
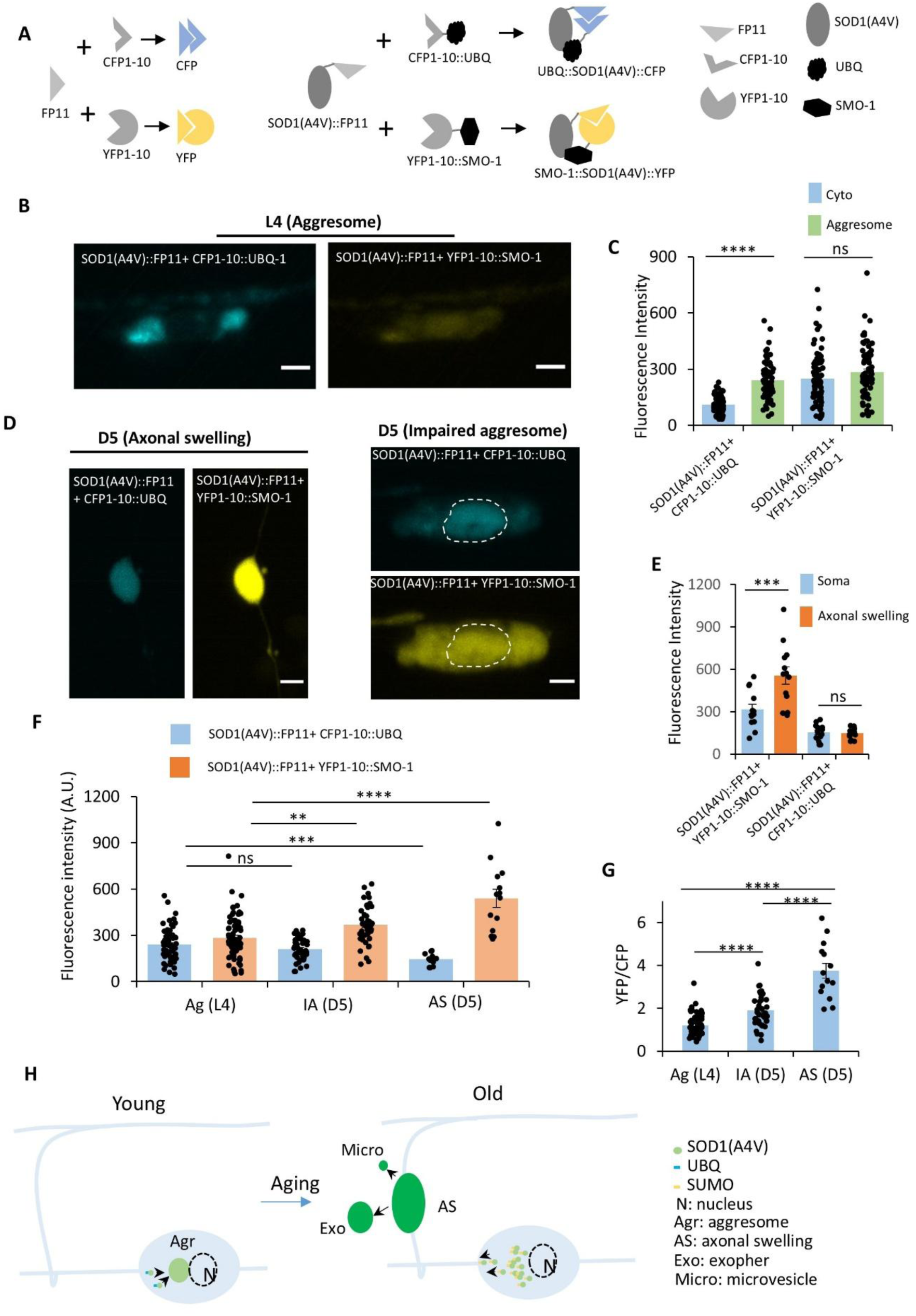
The ubiquitin-SUMO balance regulates sequestration of misfolded proteins into aggresomes and axonal swellings during aging. **A.** Schematic of the split CFP-YFP complementation technique used to detect SOD1(A4V) ubiquitination and SUMOylation. FP11 is a segment that, when bound to CFP1-10, emits CFP fluorescence and, when bound to YFP1-10, emits YFP fluorescence. FP11 is fused with SOD1(A4V), while CFP1-10 and YFP1-10 are fused with ubiquitin (UBQ-1) and SUMO (SMO-1), respectively. When SOD1(A4V) is ubiquitinated or SUMOylated, FP11 binds to CFP1-10 or YFP1-10, respectively, resulting in the emission of either CFP or YFP. **B.** Imaging of SOD1(A4V) ubiquitination using split CFP in young L4-stage animals. CFP signals are significantly brighter in the aggresomes compared to the cytoplasm. YFP indicates SOD1(A4V) SUMOylation. No YFP accumulates in aggresomes. Scale bar = 2 µm. **C.** Quantification of CFP intensity in the cytoplasm and aggresome of young L4 animals. Data are represented as mean ± SEM, n = 72, Cyto CFP vs. Aggresome CFP (P<0.0001), Cyto YFP vs. Aggresome YFP (P=0.2441), ****P < 0.0001 from one-way ANOVA with Tukey’s post hoc test. **D.** Imaging of SOD1(A4V) SUMOylation at D5 using split YFP. Left panel indicates the axonal swelling, while the right panel is the cell soma. Dashed lines in right panels outline the nuclei. Scale bar = 2 µm. **E.** Quantification of recombinant YFP and CFP intensities in the soma and axonal swellings. YFP indicates SOD1(A4V) SUMOylation, and CFP indicates SOD1(A4V) ubiquitination. N = 13, Soma YFP vs. AS YFP (P=0.0001665), Soma CFP vs. AS CFP (P=0.9987), ***P < 0.001 from one-way ANOVA with Tukey’s post hoc test. **F**-**G**. Quantification of CFP, YFP signals (**F**), and YFP/CFP ratio (**G**) in aggresomes, impaired aggresomes, and axonal swellings during ageing. Data are represented as mean ± SEM, n = 72 (L4 aggresome), 43 (D5 IA), 14 (D5 AS). Statistical comparisons in **F**: Aggresome CFP (L4) vs. IA CFP (D5) (P=0.1728), Aggresome CFP (L4) vs. AS CFP (D5) (P= 0.008124), Aggresome YFP (L4) vs. IA YFP (D5) (P= 0.0007068), Aggresome YFP (L4) vs. AS YFP (D5) (P<0.0001); Statistical comparisons in **G**: P<0.0001 for all comparisons. **P < 0.01, ***P < 0.001, ****P < 0.0001 from one-way ANOVA with Tukey’s post hoc test. **H**. Schematic of hierarchical sequestration of misfolded proteins into aggresomes and axonal swellings during aging. At young stage, misfolded proteins are tagged with ubiquitin (UBQ) and sequestered into perinuclear aggresome compartments. As animal ages, decreased UBQ and increased SUMO tagging directs misfolded proteins to axonal swellings (SolAS), where they are extruded by exophers and microvesicles. Data are represented as mean ± SEM in **C**, **E-G**.

We found that at young stages (L4), CFP but not YFP intensity was significantly higher in aggresomes than in the cytoplasm (Fig. 8B, C), indicating SOD1(A4V) ubiquitination in aggresomes. In contrast, in aged animals (D5), YFP but not CFP intensity was significantly higher in SolAS than in the cytoplasm (soma), indicating SOD1(A4V) SUMOylation within SolAS (Fig. 8D, E). These results confirm different SOD1(A4V) modifications occurring during aging.

We further compared CFP and YFP signals across different stages in aggresomes, impaired aggresomes, and SolAS. In young animals (L4), CFP was prominent in aggresomes at young age but decreased in impaired aggresomes and SolAS at D5. In contrast, YFP increased with age, peaking in SolAS at D5 stage (Fig. 8F). The YFP/CFP ratio increased with age, indicating a shift from ubiquitination-dominated to SUMOylation-dominated modifications on SOD1(A4V). This shift correlates with aggresome impairment and SolAS formation, suggesting that the balance between ubiquitination and SUMOylation regulates the hierarchical sequestration of misfolded proteins into aggresomes or SolAS during aging (Fig. 8G).

## Discussion

Our study identifies SolAS as a neuron-specific compartment that sequesters soluble misfolded proteins and expels them via exophers and microvesicles. During aging, SolAS and aggresomes function hierarchically to dispose of misfolded proteins. In young animals, ubiquitin-tagged misfolded proteins are directed to soma aggresomes. As aging progresses, the decrease in ubiquitination and aggresome function leads to SUMO-dependent SolAS formation, which reroutes misfolded proteins into neurite compartments for sequestration and removal (Fig. 8H).

Axonal swellings are often associated with neuronal injury and degeneration^15,28^, but the term appears overused to describe various types of swellings characteristic of diverse contexts. For instance, axonal swellings have been described and referred to during development, neurodegenerative diseases, and traumatic brain injury. However, they differ significantly in shape, size, cause, content, possible function, and their relationship to degeneration^15,28,37^. The SolAS we identified represents a special type of axonal swelling with a distinctive shape, selective contents, specific location, and designated function, distinct from death-related swellings. SolAS in *C. elegans* GABAergic motor neurons are elliptically shaped structures that resemble axonal torpedoes, a distinct class of axonal swellings that are not associated with neuronal death and whose function remains unclear in GABAergic Purkinje neurons^37,38^. SolAS are clearly different from the beading-like swellings that are associated with degeneration and death (Fig. S3C). SolAS form at specific segments of neurites (axonal commissures) and are closely associated with engulfing cells such as the hypodermis, unlike death-associated beading-like swellings, which form randomly along neurites (Fig. S3C). Functionally, SolAS selectively sequester soluble misfolded proteins such as polyPA and SOD1(A4V), while excluding solid aggregates like polyQ (Fig. 2N). Notably, we show that there is no microtubule looping or axonal degeneration before or after SolAS-mediated expulsion, distinguishing these structures from pathological axonal swellings with microtubule looping that blocks axonal transport and causes degeneration^28^. These findings suggest that SolAS represent a distinct class of axonal swellings designed to manage soluble misfolded proteins without significantly compromising neuronal function.

Quality control compartments are protective structures because they facilitate the degradation of misfolded proteins. Aggresomes and ER-associated compartments (ERAC) employ autophagy to degrade misfolded proteins, thereby reducing their intracellular levels^1,3,39,40^. Our findings suggest that, instead of degrading intracellularly, SolAS utilize a distinct mechanism of extracellular extrusion, to reduce misfolded protein levels, via exophers or microvesicles. Exophers were first identified in *C. elegans* touch receptor neurons as a mechanism for expelling neuronal waste from the soma^26^. Interestingly, this mechanism is employed by SolAS in neurites to reduce misfolded protein levels in GABAergic motor neurons. Notably, both the somata of touch receptor neurons and commissures of the GABAergic neurons lie adjacent to the hypodermis^21,22^, indicating that anatomical positioning facilitates this clearance process.

The formation of aggresomes and SolAS depends on different tagging molecules. Misfolded proteins are tagged with ubiquitin for sequestration into the aggresome^41^. Our study shows that ubiquitin tagging also contributes to maintaining aggresome integrity in young animals. However, as aging progresses, global ubiquitination gradually declines^31^, which may reduce aggresome integrity, causing the dispersion of misfolded proteins within neurons. Unlike ubiquitination, global SUMOylation increases with age in *C. elegans*^32^. These misfolded proteins are then tagged with SUMO and transported to the commissure, where they are sequestered into SolAS. Our results suggest a dynamic ubiquitin-SUMO balance in regulating quality control compartments. In young animals, misfolded proteins are tagged with ubiquitin and sequestered into aggresomes, where they are processed and degraded through autophagy. However, as ubiquitin function declines with age, misfolded proteins are instead tagged with SUMO, sequestered into SolAS, and extruded via exophers and microvesicles.

## Methods

### *C. elegans* Strains and Maintenance

*C. elegans* were maintained at 20 °C on Nematode Growth Medium (NGM) agar plates seeded with *E. coli* OP50, following standard protocols^42^. Strains used in this study were listed in the Supplemental table S1 strain list.

### Molecular Biology and Cloning

We used the pBS77 vector as the cloning backbone. Neuron-specific promoters, such as *Punc-25* (D-type motor neuron), *Punc-129* (DA/DB motor neuron), or *Pacr-15* (SIA neuron), were inserted into the vector using SbfI and BamHI restriction sites. Human SOD1 was obtained by PCR amplification from the genomic DNA of an hSOD1 integrated worm strain AM263 rmIs175 [*unc-54p::hSOD1 (WT)::YFP*]^43^ obtained from the *Caenorhabditis* Genetics Center and subsequently inserted into a vector with *Punc-25* promoter via KpnI and NotI sites. *hSOD1(A4V), hSOD1(G93), hSOD1(G147P),* and *hSOD1(N53I)* mutations were introduced by PCR-based site-directed mutagenesis using *hSOD1 (WT)::YFP* plasmid as a template. To generate the split CFP-YFP complementation system for monitoring ubiquitin-and SUMO-modifications on SOD1(A4V), we fused FP11 with SOD1(A4V) (C-terminal), split CFP (1-10) with ubiquitin (N-terminal), and YFP (1-10) with SUMO (N-terminal). GA50 and PA50 were obtained from Addgene^44^ and inserted into pBS77 using KpnI and AscI. All markers, including *tbg-1, sqst-1, ubq-1, smo-1, rpn-5, ifp-1, lmp-1*, and *lgg-1* were cloned from *C. elegans* genomic DNA. All primers were listed in Supplemental table S2 primer list.

CRISPR-Cas9 genome editing was carried out as previously described^45^. The mix for microinjection included 5 μM sgRNA, 5 μM Cas9 nuclease, and a single-stranded oligonucleotide (ssODN) repair template, injected into the gonads. Three independent strains with high transmission rates were genotyped to verify the mutations.

### Microinjection and Integration

Microinjections were performed using an inverted Olympus CX51 microscope equipped with an Eppendorf FemtoJet 4i. Day 1 young adult worms were selected and immobilized on a 2% agarose pad and covered with a drop of halocarbon oil for microinjection. Plasmid concentrations ranging from 1 ng/μL to 25 ng/μL were injected into *C. elegans* gonads. Transgenic strains with high transmission rates of the extrachromosomal arrays were selected for subsequent experiments.

Some of the extrachromosomal arrays were integrated into *C. elegans* genome using TMP-UV method. 70 worms carrying extrachromosomal arrays were treated with 30 μg/mL TMP for 15 minutes. Worms were then subjected to UVC radiation using a UV crosslinker. TMP-UV-treated P0 worms were singulated (one worm per plate) and allowed to propagate to the F2 generation. Approximately 200 F2 progeny were selected to screen for successful genomic integrations. Positive integrated strains were backcrossed with N2 wild-type worms five times to remove background mutations.

### Confocal imaging

Confocal imaging experiments were conducted using a spinning-disk confocal microscope (Diskovery, Andor Technology) at the Advanced Microscopy Facility of Queensland Brain Institute. Prior to imaging, worms were transferred to an NGM plate and paralysed with a drop of 5 mM tetramisole hydrochloride. Three to five paralyzed worms were then transferred into a small drop of 5 mM tetramisole hydrochloride on an agarose pad (6%). Because the commissures of D-type motor neurons are on the right side of the worm body, mispositioning could result in blurry neurite images and impede neuron tracing. To obtain stable and clear images, we carefully adjusted the worm position to let its right side face up before placing a cover glass. This orientation ensures that commissures of D-type motor neurons face the objective directly on the inverted confocal microscope. Worms were then subjected to 3D imaging on the confocal microscope with a Z-step of 0.15 μm. Images were captured in segments from tail to head, then stitched together to form a complete whole-body pattern of D-type neurons. Image stacks were projected using maximum intensity in ImageJ before analysis. For all SOD1::YFP and PA50::GFP imaging during aging, the same imaging conditions, including exposure time and laser power, were used to allow consistent comparisons across different ages and animals.

The stitched whole-body images were used for neuron identification and neurite tracing. The 19 D-type motor neurons on the ventral side were divided into six groups (from head to tail, group 1 to group 6, Fig. S1A). The first five groups (group 1 to group 5) each have two VD neurons and one DD neuron (VD1 to VD10 and DD1 to DD5), the sixth group has three VD neurons and one DD neuron (VD11, VD12, VD13 and DD6). Small gaps on the ventral side were often observed between different groups of neurons. As commissures in group 1 neurons were difficult to separate, we only quantified neurons in groups 2 to group 6, in total 16 neurons (DD2-DD6 and VD3-VD13) in our experiments. Identification and tracking started from the tail end and use the neuron VD11 as a landmark to separate the fifth and sixth groups of neurons (this neuron has relatively long distance from both the fifth group of neurons and the remaining three neurons in the sixth group. After identifying these neurons, commissural neurites from groups two to six were traced back to their respective somata.

After ID each of the neurons in group 2 to group 6, we quantified compartment structures in these neurons. For aggresome identification, we used the following criteria: 1) perinuclear structures with higher fluorescent intensity than surrounding areas and obvious boundaries; 2) typically one to two events within a single neuron. For SolAS identification, we used the following criteria: 1) spindle-shaped structures packed with accumulating misfolded proteins in commissures; 2) usually one per neuron. Exophers were identified as larger vesicles, 1–5 μm in size, often located near or connected to SolAS by a thin tube structure. Microvesicles were classified as extracellular vesicles, 100–1000 nm in diameter, located near or budding from axonal swellings.

We used longitudinal imaging to track the same aggregates within the same neuron in the same animals over the course of aging. The longitudinal imaging was achieved by multiple rounds of imaging-recovery cycles at different ages. Ten to twenty L1 worms were singulated to different numbered plates. To minimize potential damage during imaging and mounting, we only placed one worm on a single agarose pad at a time. After imaging, the animal was carefully and quickly transferred to a NGM plate with OP50 for recovery. These imaging-recovery cycles were repeated for the same animal every other day until the worm reached D11 (we stopped at D11 as worms after this stage are very fragile). Not all worms reach D11 due to damage or loss during the recovery process. Worms with at least D7 data were included in our analysis.

These longitudinal imaging data were then used for the Hidden Markov Model (HMM) analysis. Group two to six neurons with their neurites were first annotated according to Fig. S1A. Each individual neuron was then assigned one of four states: 1. Diffuse state (D): SOD1(A4V)::YFP is diffusely distributed throughout the neuron, with no visible aggregates in the soma or neurites. 2. Aggresome state (Agr): Aggresomes form in the soma, but no axonal swellings are observed in the neurites. 3. Impaired Aggresome state (IA): Aggresomes lose integrity or become undetectable, with no axonal swellings present in the neurites. 4. Axonal Swelling state (AS): Soma aggresomes are impaired, and axonal swellings are observed at the commissure. A state map was created by linking each neuron states at different ages for each individual animal. To determine overall state transition probabilities, all transitions originating from identical starting states were pooled together and treated as one. Each transition was then quantified by the percentage of transitions into different states. Transition pathways were constructed by connecting state transitions with transition probability values.

### FRAP and Photoactivation

Fluorescence recovery after photobleaching (FRAP) and photoactivation of PA-GFP were conducted using a Zeiss LSM 710 confocal microscope with Zeiss Zen Black software. For FRAP, L4 or D3 animals were immobilized and placed on a 6% agarose pad. Time-lapse images were acquired with 488 nm lasers at low power (0.11%) for approximately 1 minute. Photobleaching at the selected region of interest (ROI) was conducted with a high-powered 488 nm laser beam (100%) for five cycles, starting at the 2^nd^ second. Mean grey values from the bleached area were calculated and normalized based on values from a distant, unbleached region (we used fluorescent signal in nucleus).

For photoactivation, ten to twenty L4 stage worms were isolated into different plates. As these experiments require multiple rounds of imaging on the same animal, we placed only one worm on a single agarose pad at a time to minimize potential damage during imaging and mounting. Imaging was initiated with L4-stage worms immobilized and mounted on a 6% agarose pad. To minimize phototoxicity and ensure good recovery, we only photoactivated one neuron in one worm. Neurons with prominent aggresome structures (using RPN-5::mKate2 as aggresome marker) were selected for photoactivation. Pre-activation images were captured using low power of 488 nm and 561 nm light. Photoactivation of PA-GFP was achieved using 15% power of 405 nm light, followed by post-activation imaging with low power of 488 nm and 561 nm light. Worms were quickly recovered and transferred to fresh NGM plates with OP50 for three-day maintenance, and the same neuron was re-imaged at D3 with low power of 488 nm and 561 nm light to check SolAS signal. Image analysis was performed using ImageJ.

### Drug Treatment

Nocodazole (Sigma Aldrich, USA) was used to disrupt microtubules^46^ in *C. elegans* neurons. NGM plates were supplemented with Nocodazole to a final concentration of 5 µM and seeded with OP50. Synchronized L4-stage worms were transferred to drug-positive or control (DMSO) plates and allowed to grow for three days. D3 worms were then immobilized and mounted on a 6% agarose pad for imaging experiments. Latrunculin A (LatA, Novachem) was used to inhibit actin polymerization^47^. L4-stage worms were transferred to NGM plates containing LatA at a final concentration of 4 μM or to a DMSO control plate. Worms were grown to D3 and mounted for imaging experiments.

### Lifespan assay

Lifespan assays were conducted as previously described^48^. 6 cm NGM plates used for lifespan assays were supplemented with a final concentration of 50 µM 5-fluorodeoxyuridine (FUDR) and seeded with OP50. 100 L4-stage worms were transferred to 10 plates. Worms were transferred every three to four days until all had died. Survival was assessed by gently touching the tail region; no response indicated death, and such worms were removed from the plates. Worms that crawled off the plates or burst were censored from the analysis. Survival analysis was performed using the Online Application for Survival Analysis 2 (OASIS 2)^49^ (https://sbi.postech.ac.kr/oasis2/). The log-rank method was employed for the significance test.

### Statistics and reproducibility

We used online OASIS2 for lifespan analysis and statistics, and used GraphPad Prism 10 for all other statistics in this study. Paired or unpaired Student’s t-tests were used for two-group comparisons, while one-way or two-way ANOVA was employed for multiple group comparisons. The log-rank test was used for lifespan comparisons. Sample sizes, p-values, and statistical tests were indicated in figure legends. Data were presented as the mean value ± SEM, and p < 0.05 was deemed a significant difference. Unless indicated otherwise, graphs show all biological replicates collected across three or more independent replicates. No statistical methods were employed to determine the sample sizes; however, our sample sizes are comparable to those used in previous studies^50^. For imaging experiments, animals were randomly selected, excluding dead, bagging, or burst worms. In the lifespan assay, worms were censored if they crawled off the plates or burst.

## Data availability

The data from this study is available in the source data file. Further information and requests for resources and reagents are available from the lead contact upon request.

## Acknowledgements

We thank the *Caenorhabditis* Genetics Center (CGC) and National BioResource Project (NBRP) for providing *C. elegans* strains, and Addgene for providing plasmids. We thank Rumelo Amor and other members of QBI Advanced Microscopy Facility for technical assistance. We thank Prof Pankaj Sah, A/Prof Zhitao Hu and Dr Sean Coakley for invaluable discussions. We thank Dr Brent Neumann for his thoughtful suggestions on the manuscript. This work was supported by NHMRC IDEAS Grants (GNT2002472, to Z.L.), Bartlett Senior research fellowship and Queensland Brain Institute start-up funding (to Z.L.).

## Author contributions

Conceptualization: Z.L. Study design: M.P.L., Y.W., and Z.L. Experimentation: M.P.L and Y.W. carried out most experiments. Y.R., Y.C., X.Y., M.Z., H.L., Y.Y., Y.C., Y.Y. and Y.L. contributed to some experiments. Data analysis and interpretation: M.P.L., Y.W., Y.R., Y.C., X.Y., M.Z., H.L., Y.Y., Y.C., K.X., Y.Y., D.S., Y.L., V.A., M.A.H., and Z.L. Visualization: M.P.L and Y.W., M.Z., H.L., and Z.L. Supervision: Z.L. Writing-original draft: M.P.L., Y.W., and Z.L. Writing-review & editing: M.P.L., Y.W., V.A., M.A.H., and Z.L.

## Supplemental information

Document S1.

---

**Fig. S1:**
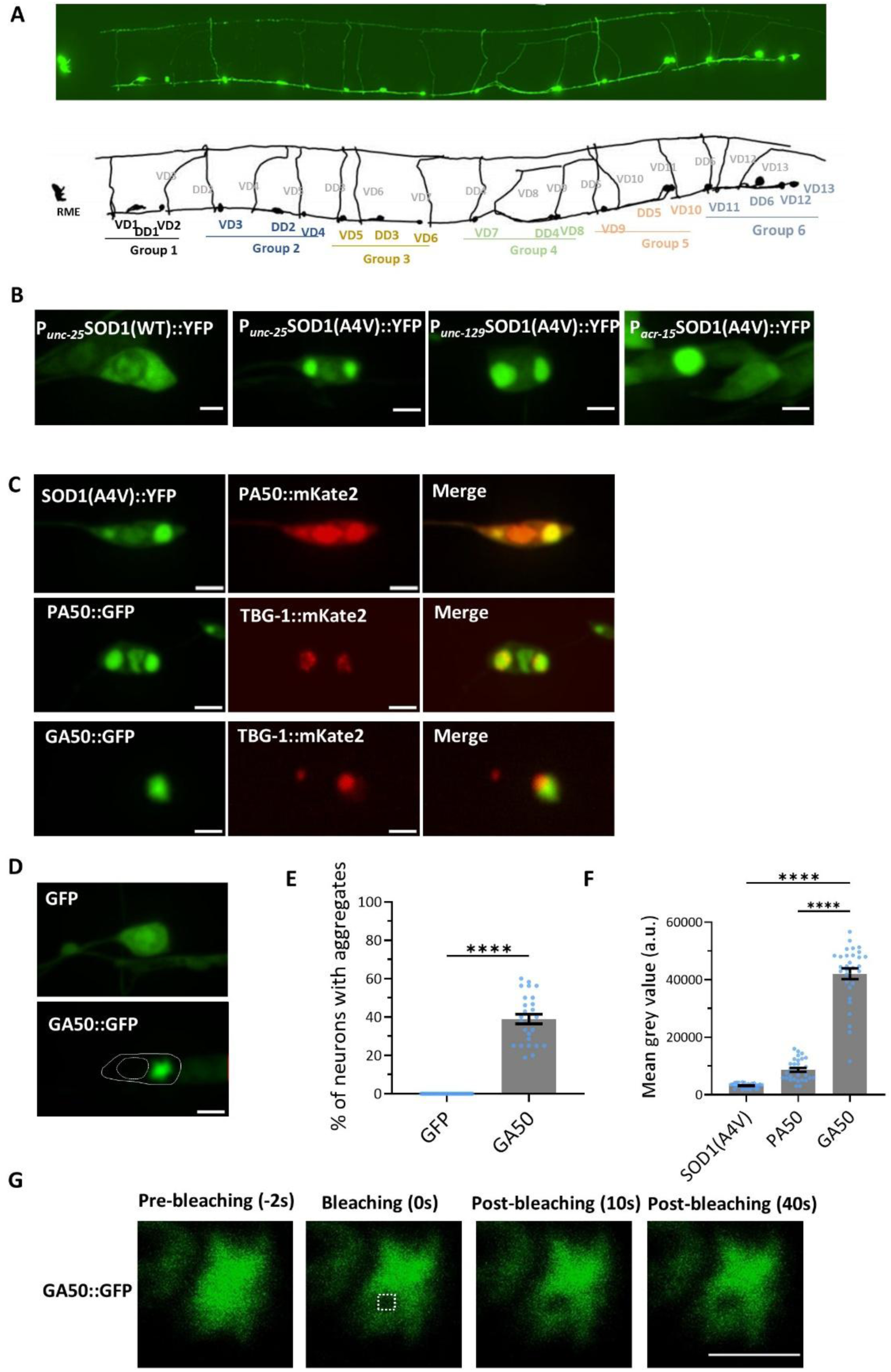
Some neurodegenerative disease-associated misfolded proteins are sequestered into aggresomes in *C. elegans* neurons. **A.** Organization of D-type motor neurons. 19 D-type motor neurons on the ventral side are divided into six groups: the first five groups each contain three neurons (two VD neurons and one DD neuron), and the sixth group consists of four neurons (three VD neurons and one DD neuron). Commissure neurites from groups 2 to 6 can be traced back to their respective soma, while commissure neurites in group 1 often overlap, making them difficult to distinguish. In all experiments, only group 2 to group 6 neurons were quantified. The *Punc-25* promoter was used to drive expression in D-type GABAergic motor neurons. **B.** Expression of SOD1(A4V) in different types of neurons triggers the formation of similar aggregates. The *Punc-25* promoter drives expression in D-type GABAergic motor neurons, *Punc-129* promoter drives expression in cholinergic motor neurons, and the *Pacr-15* promoter drives expression in sublateral motor neurons SIA. **C.** Colocalization of PA50 and GA50 with the aggresome marker TBG-1. PA50 aggregates, but not GA50 aggregates, strongly colocalize with TBG-1. Scale bar = 2 µm. **D.** GA50 expression in *C. elegans* GABAergic motor neurons forms fibrillar amyloid-like aggregates. The solid line outlines the neuron body, and the dashed line outlines the nucleus. Scale bar = 2 µm. **E.** Quantification of GA50 aggregate formation in GABAergic motor neurons. Data are presented as mean ± SEM, n = 26, ***P < 0.001, determined by unpaired t-test. **F.** Quantification of fluorescence intensity in the aggregate regions of SOD1(A4V), PA50, and GA50. Data are presented as mean ± SEM, n ≥ 30, ****P < 0.0001 from one-way ANOVA with Tukey’s post hoc test. **G.** Photobleaching of GA50::GFP. Images were presented at −2 s, 0 s (immediately before photobleaching), 10 s, and 40 s post-bleaching. Dash box indicates photobleached region. Photobleaching was performed at 0 s. GA50::GFP exhibited solid-like properties. Scale bar = 2 µm.

**Fig. S2:**
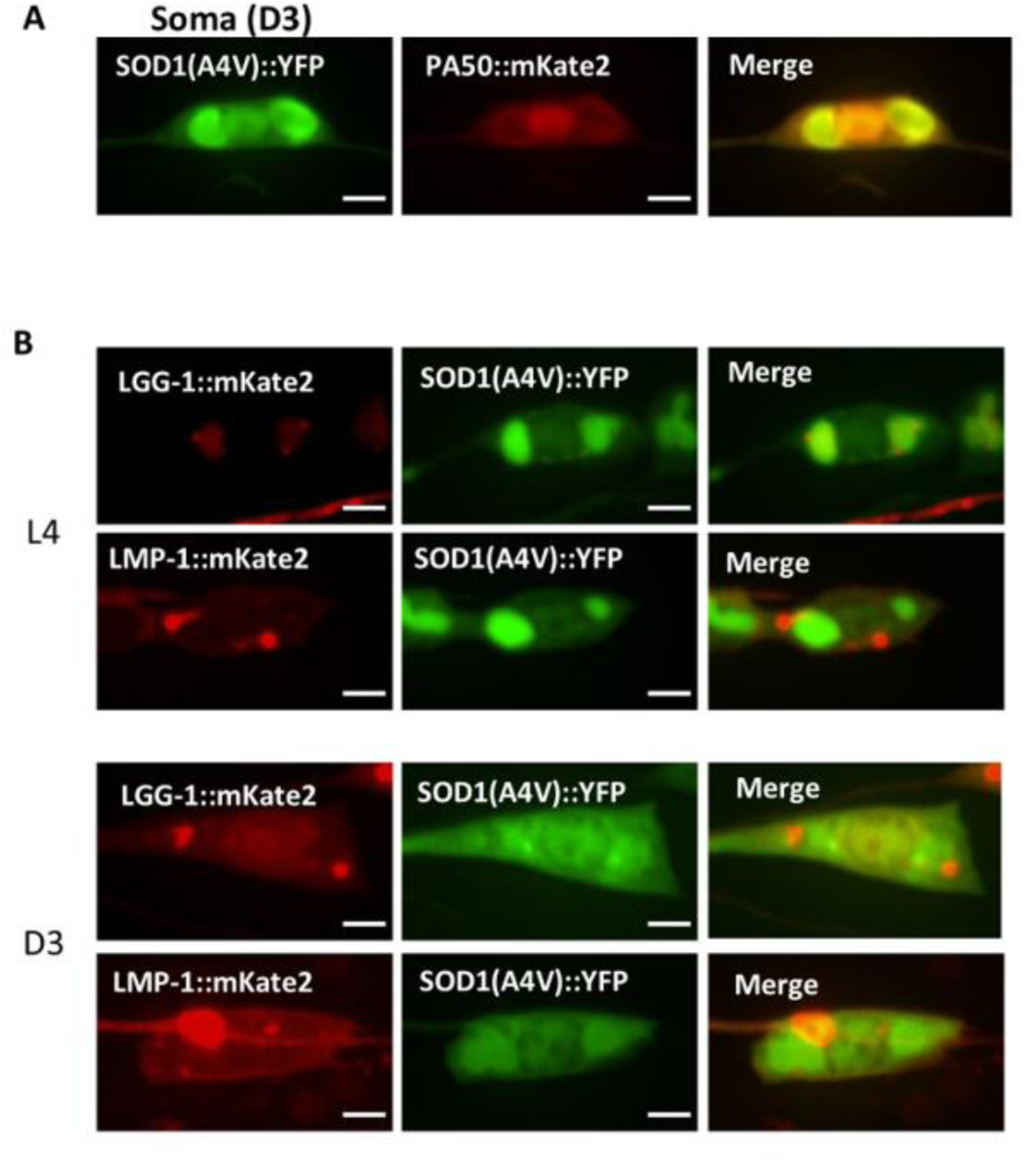
Misfolded protein associated impaired aggresomes are not fully colocalized with autophagy and lysosome markers. **A.** Colocalization of SOD1(A4V)::YFP with PA50::mKate2 in the neuron soma at the D3 stage. Scale bar = 2µm. **B.** Aggresomes do not fully colocalize with the autophagosome marker LGG-1 or lysosome marker LMP-1 before and after aggresome impairment. LGG-1 puncta are observed within or surrounding the aggresomes, and some LMP-1 puncta are found surrounding or near aggresomes. Scale bars = 2µm.

**Fig. S3:**
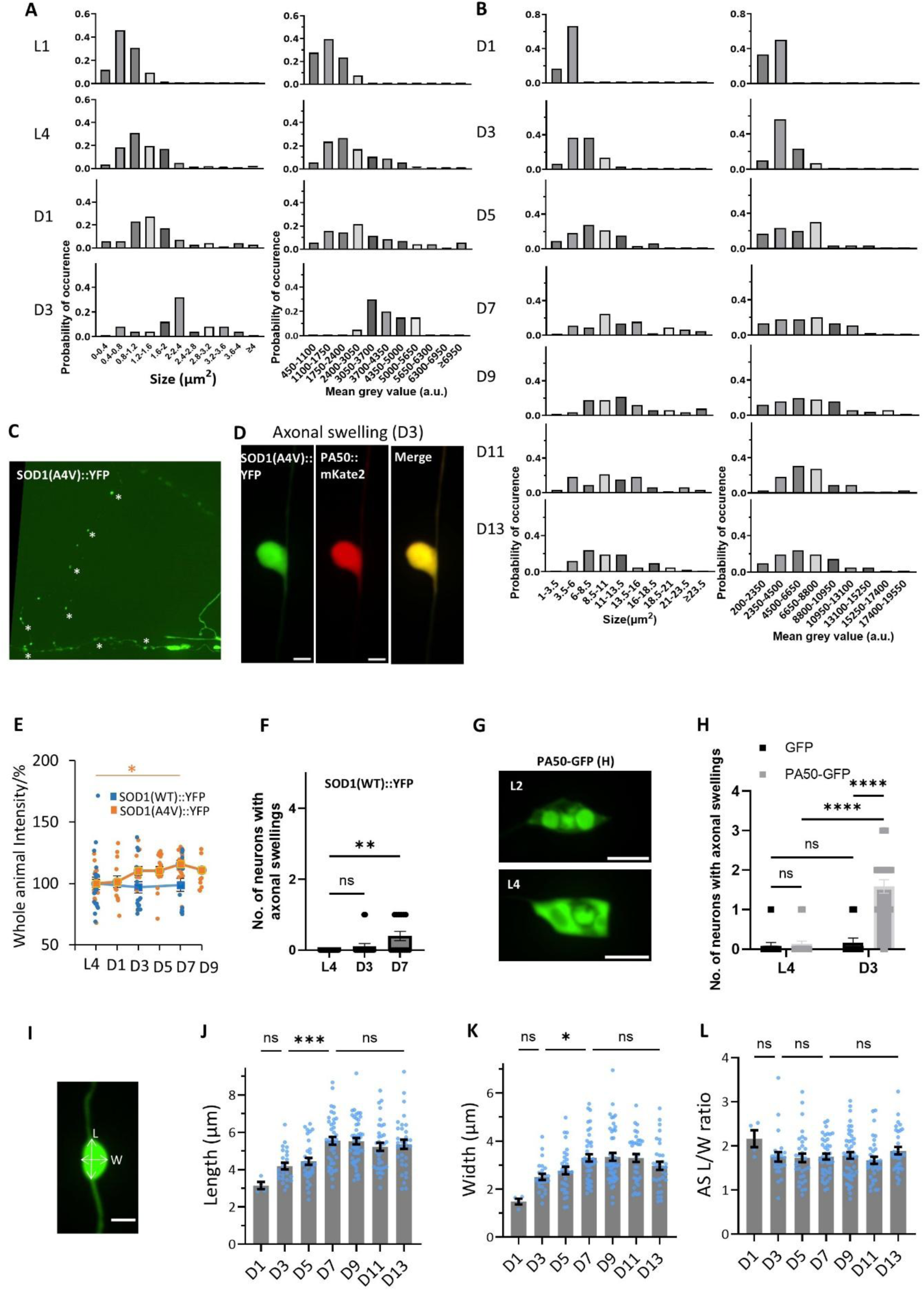
Misfolded proteins are sequestered into aggresomes and axonal swellings during aging. **A.** Histograms showing the size (left panel) and intensity (right panel) of aggresomes during aging. Data is presented from the L1 to D3 stages, as aggresomes are rare after D3. N = 457. **B.** Histograms showing the size (left panel) and intensity (right panel) of axonal swellings during aging. The L4 stage is excluded due to the absence of axonal swelling events. N = 232. **C.** Formation of beading-like swelling along the neurite prior to neuron death caused by long-time sodium azide treatment in SOD1(A4V)::YFP worms. Asterisks indicate beadings along the neurite. **D.** Colocalization of SOD1(A4V)::YFP with PA50::mKate2 in axonal swellings at the D3 stage. Scale bar = 2 µm. **E.** Quantification of whole worm (in all the D-type motor neurons) SOD1(WT) and SOD1(A4V) levels during aging. SOD1(WT): n = 21(L4), 18(D3), 15(D7); SOD1(A4V): n = 16 (L4), 13 (D1), 15 (D3), 17 (D5), 17 (D7), 16 (D9), *P = 0.01582 (SOD1(A4V) L4 vs. D7), one-way ANOVA with Tukey’s post hoc test. **F.** Quantification of axonal swelling numbers in neurons expressing SOD1(WT) during aging. N = 21(L4), 18(D3), 15(D7), p = 0.4729 (ns), **p = 0.0012, one-way ANOVA with Tukey’s post hoc test. **G.** Impairment of aggresomes at L4 stage with high level of PA50 expression in DD neuron. Scale bar = 4 µm. **H.** Quantification of axonal swelling numbers in DD neurons expressing high levels of PA50 during aging. L4 stage: n = 12(GFP), 23 (PA50-GFP); D3 stage: n = 12 (GFP), 24 (PA50-GFP). P > 0.9999 (L4 GFP vs. L4 PA50-GFP), P = 0.995 (L4 GFP vs. D3 GFP), ****P < 0.0001 (L4 PA50-GFP vs. D3 PA50-GFP; D3 GFP vs. D3 PA50-GFP), two-way ANOVA with Šídák’s multiple comparisons test. **I.** Example showing the method for measuring the length and width of a torpedo-like axonal swelling. **J**-**L**. Quantification of axonal swelling length (**J**), width (**K**), and the length/width ratio (**L**) during aging. As both length and width increase, no significant change in the length/width ratio was observed during aging. Data are presented as mean ± SEM, P=0.6904, 0.0004, 0.994, 0.4474, 0.0281, 0.7739, 0.7265, 0.9999, 0.9068 from **J** to **L**, *p < 0.05, ***p < 0.001, one-way ANOVA with Tukey’s post hoc test. Data are represented as mean ± SEM in **E-F, H, J-L**.

**Fig. S4:**
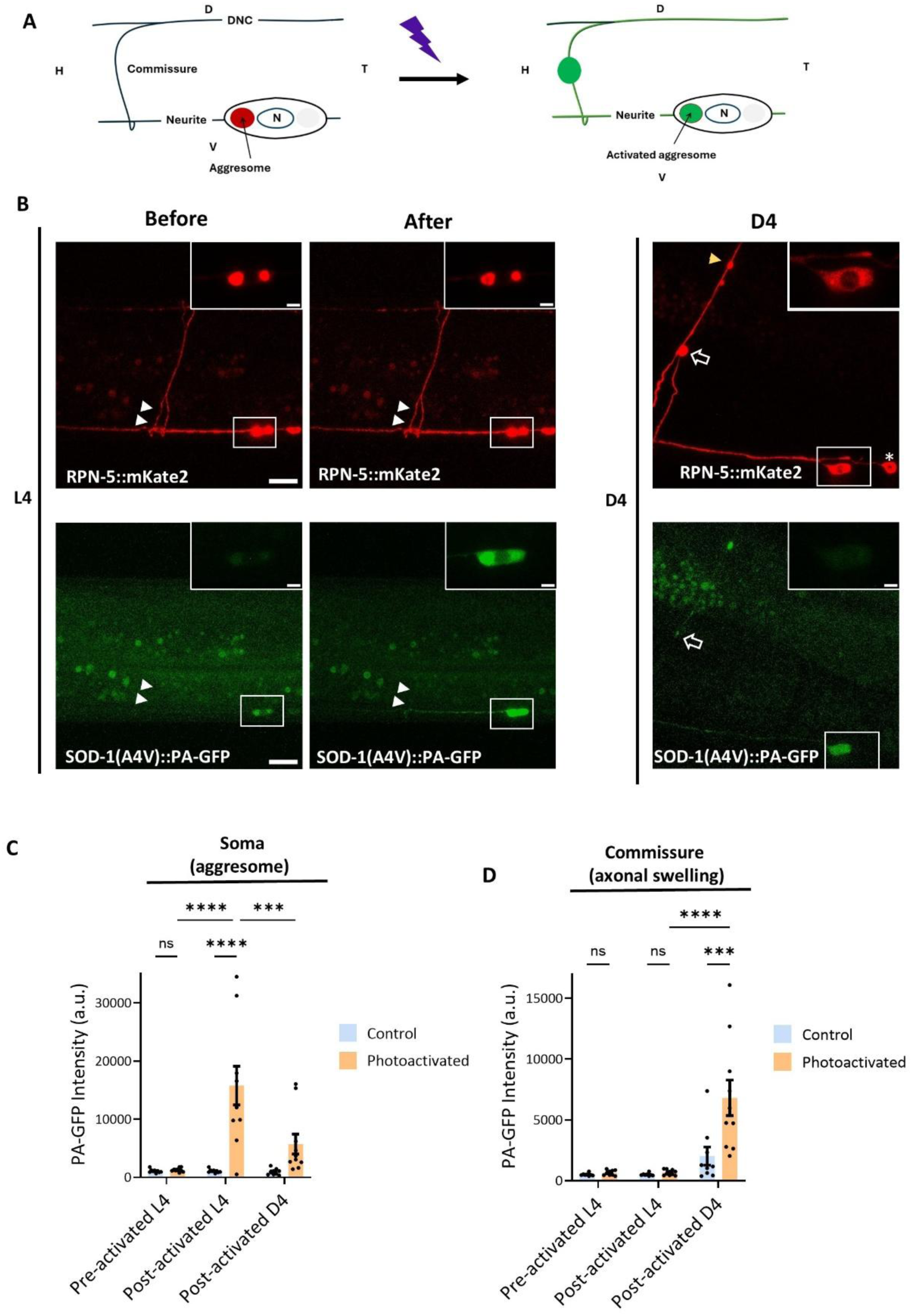
Misfolded proteins are transported from impaired aggresomes to axonal swellings during aging. **A.** Schematic of the photoactivation experiment. SOD1(A4V)::PA-GFP was expressed in *C. elegans* GABAergic neurons. At the L4 stage, when aggresomes form, photoactivation was performed by activating PA-GFP at the aggresome using 405 nm light. Imaging was conducted both before and after photoactivation. Worms were then recovered and maintained in a dark environment. At D4, when axonal swellings formed, imaging was performed again. D-dorsal, V-ventral, H-head, T-tail, N-nucleus. **B.** Changes in PA-GFP localization in neurons before and after photoactivation. RPN-5::mKate2 was used as an aggresome marker. White arrows indicate commissures (L4), while yellow arrows indicate axonal swellings (D4) of the same neuron (white box) after photoactivation. The hollow arrow indicates an axonal swelling in a neighbouring neuron (marked with an asterisk) that was not photoactivated. Note that PA-GFP in the aggresome shows increased brightness after photoactivation at L4, and bright fluorescence is observed in the axonal swelling at D4. Fluorescence intensity in the axonal swelling of the neuron without photoactivation remains low. Scale bar = 5 µm and 1 µm (zoom in window). **C.** Quantification of PA-GFP intensity in the soma before and after photoactivation. Data are presented as mean ± SEM, n ≥ 9, P > 0.9999, < 0.0001, < 0.0001, = 0.0007 from left to right, ***p < 0.001, ****p < 0.0001, one-way ANOVA with Tukey’s post hoc test. **D.** Quantification of PA-GFP intensity before and after photoactivation at the commissure and axonal swelling. Data are presented as mean ± SEM, n ≥ 9, P > 0.9999, < 0.0001, < 0.0001, = 0.0003 from left to righg, ***p < 0.001, ****p < 0.0001, one-way ANOVA with Tukey’s post hoc test.

**Fig. S5:**
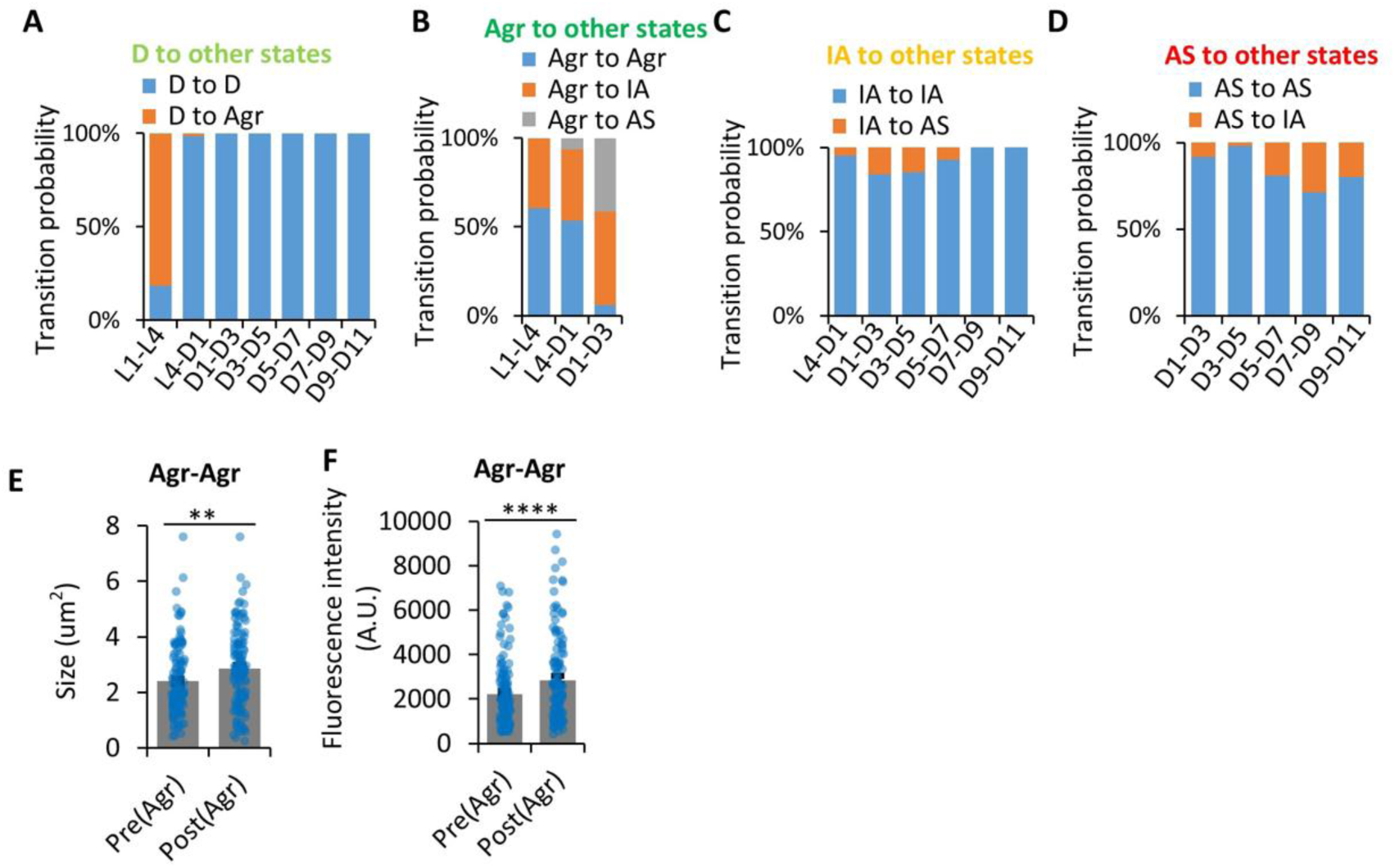
Hierarchical sequestration of toxic proteins into aggresomes and axonal swellings during aging. **A.** Transition probability from the disperse state (D state) to other states (D state and Agr state) during aging. n = 70 (L1-L4), n = 61 (L4-D1), n = 53 (D1-D3), n = 50 (D3-D5), n = 40 (D5-D7), n = 26 (D7-D9), n = 5 (D9-D11). **B.** Transition probability from the aggresome state (Agr state) to other states (Agr state, IA state, and AS state) during aging.n= 89 (L1-L4), n = 125 (L4-D1), n = 65 (D1-D3) **C.** Transition probability from the impaired aggresome state (IA state) to other states (IA state and AS state) during aging. n = 43 (L4-D1), n = 82 (D1-D3), n = 104 (D3-D5), n = 85 (D5-D7), n = 46 (D7-D9), n = 9 (D9-D11). **D.** Transition probability from the axonal swelling state (AS state) to other states (AS state and IA state) during aging. n = 12 (D1-D3), n = 52 (D3-D5), n = 63 (D5-D7), n = 35 (D7-D9), n = 10 (D9-D11). **E**-**F**. Quantification of aggresome size(**E**) and fluorescence intensity change (**F**) when a neuron undergoes Agr-Agr state transition. If one neuron has more than one aggresome, the size value is the sum of all the aggresomes in that neuron. Data are presented as mean ± SEM, n = 126, P=0.0019 in **E** and P<0.0001 in **F**. **P < 0.01, ****P < 0.0001 from paired t-test.

**Fig. S6.**
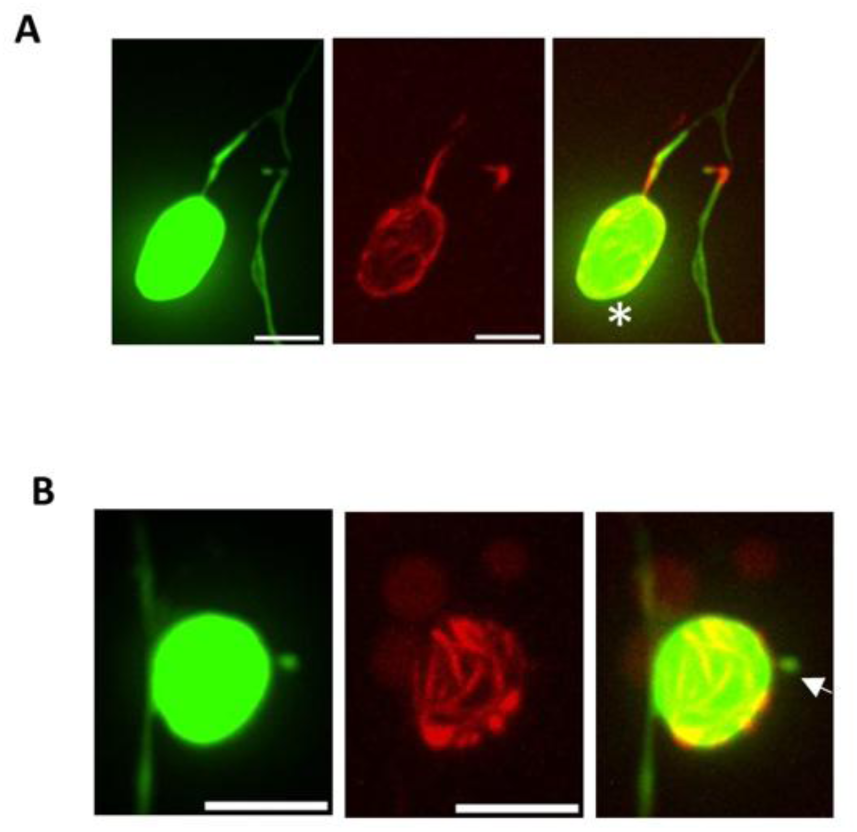
Cage-like actin network is observed in exophers but not microvesicles. **A.** A cage-like actin network colocalizes with exopher. A white asterisk indicates the exopher. Scale bar = 4 µm. **B.** Actin network does not colocalize with microvesicle. The white arrow indicates the microvesicle. Scale bar = 4 µm.

**Fig. S7:**
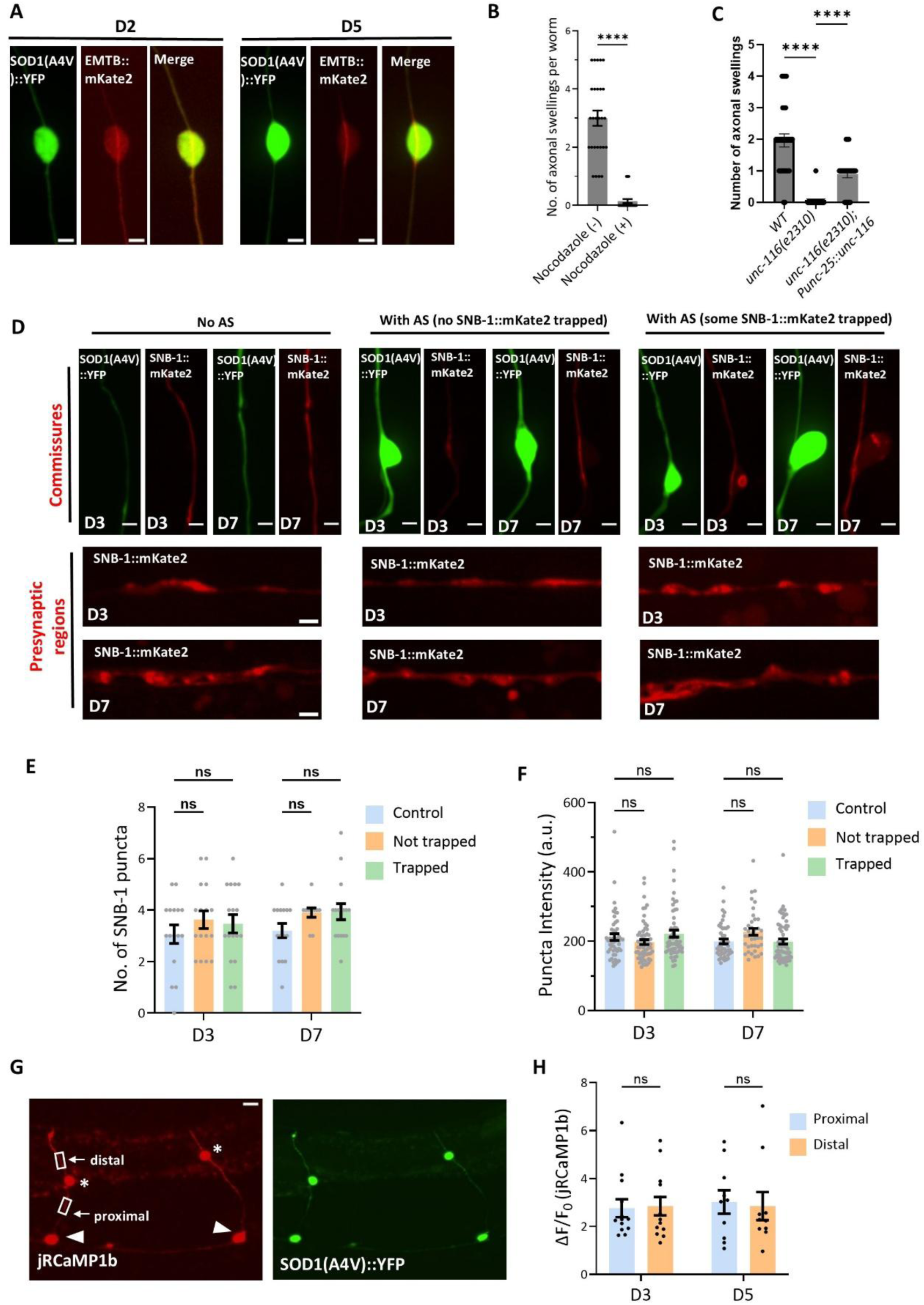
Impact of axonal swellings on microtubule function and neuronal excitability propagation. **A.** Microtubule organization remains intact within SOD1(A4V)-induced axonal swellings at D2 and D5. EMTB-labeled microtubules show no evidence of looping within these swellings. Scale bar: 2 µm. **B.** Quantification of axonal swelling numbers following treatment with nocodazole or DMSO (control). Treatment initiated at the L4 stage and imaging performed at D3. N = 27 for each group, ****p < 0.0001 from unpaired t-test. **C.** Quantification of axonal swelling numbers in kinesin mutants *unc-116* and the rescue strains. N= 29 for each group, ****p < 0.0001 from one-way ANOVA with Tukey’s post hoc test. **D.** Axonal swellings do not significantly hinder the transport of SNB-1 to presynaptic regions. Neurons are categorized based on axonal swelling presence in commissures and SNB-1::mKate2 trapping: 1) no axonal swelling (control), 2) axonal swelling without trapped SNB-1::mKate2, and 3) axonal swelling with trapped SNB-1::mKate2. SNB-1::mKate2 signals at commissure segments and dorsal side presynaptic regions are captured at D3 and D7 stages. **E.** Quantification of SNB-1 puncta across the three groups: no axonal swelling, axonal swelling without trapped SNB-1::mKate2, and axonal swelling with trapped SNB-1::mKate2. N ≥ 10, P=0.3732, 0.6242, 0.0693, 0.9993 from left to right. Dunnett’s multiple comparisons test. **F.** Quantification of presynaptic SNB-1 puncta intensity at the dorsal side across the three groups. N ≥ 39, P=0.3519, 0.5542, 0.3069, 0.1933 from left to right. Dunnett’s multiple comparisons test. **G.** Calcium imaging of axonal swellings using jRCaMP1b at distal and proximal segments of commissures. Asterisks mark axonal swellings. White boxes highlight proximal and distal segments for calcium imaging. White triangles indicate neuron soma. **H.** Quantification of calcium changes at proximal and distal segments at D3 and D5. N ≥ 10, P=0.999, 0.9939 from left to right. One-way ANOVA with Tukey’s post hoc test. Data are presented as mean ± SEM in **B-C, E-F, H**.

**Fig. S8:**
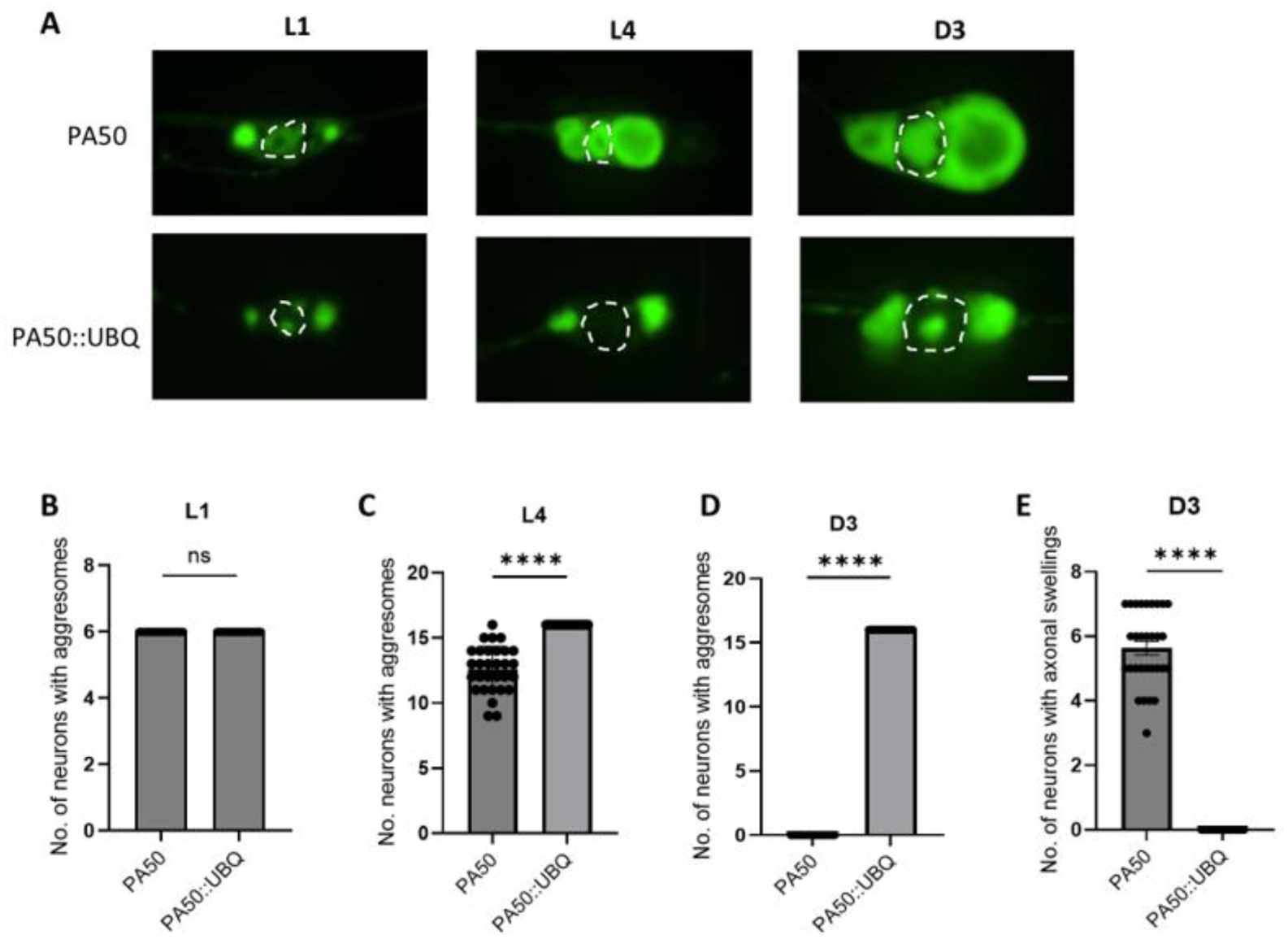
PA50 fusion with ubiquitin maintains aggresome integrity and suppresses axonal swelling in neurons during aging. **A.** Fusion of ubiquitin with PA50 prevents aggresome impairment and preserves aggresome integrity during aging. Imaging was conducted at L1, L4, and D3 stages. Dash line indicates nucleus. Scale bar = 2 µm. **B**-**D**. Quantification of neurons with aggresomes in animals with or without ubiquitin fusion to PA50 at L1 (**B**), L4 (**C**), and D3 (**D**) stages. Data are presented as mean ± SEM, n = 24 (**B**), 30 (**C**), 30 (**D**), ****p < 0.0001 from unpaired t-test. **E.** Quantification of neurons exhibiting axonal swellings in animals with or without ubiquitin fusion to PA50 at the D3 stage. Data are presented as mean ± SEM, n = 30, ****p < 0.0001 from unpaired t-test.

**Supplemental table S1:** Strain list.

|  |  |
| --- | --- |
| *C. elegans* strain: wild type | N2 |
| *atgEx5*[*Punc-25::GFP*] | DNA5 |
| *atgls20*[*Punc-25::SOD1(WT)::YFP*] | DNA8 |
| *atgls9*[*Punc-25::SOD1(A4V)::YFP*] | DNA9 |
| *atgEx10*[*Punc-25::SOD1(N53I)::YFP*] | DNA10 |
| *atgEx11*[*Punc-25::SOD1(G93A)::YFP*] | DNA11 |
| *atgEx12*[*Punc-25::SOD1(G147P)::YFP*] | DNA12 |
| *atgls253* [*Punc-25::PA50::GFP*] | DNA253 |
| *atgls603*[*Punc-25::mK2::TBG-1*]; *atgls9*[*Punc-25::SOD1(A4V)::YFP*] | DNA6003 |
| *atgEx62*[*Punc-25::VIMENTIN::mK2*]; *atgls9*[*Punc-25::SOD1(A4V)::YFP*] | DNA6002 |
| *atgEx532*[*Punc-25::mK2::IFP-1*]; *atgls9*[*Punc-25::SOD1(A4V)::YFP*] | DNA6004 |
| *atgls84*[*Punc-25::mK2::UBQ-1*]; *atgls9*[*Punc-25::SOD1(A4V)::YFP*] | DNA6005 |
| *atgls83*[*Punc-25::SQST-1::mK2*]; *atgls9*[*Punc-25::SOD1(A4V)::YFP*] | DNA6006 |
| *atgls82*[*Punc-25::RPN-5::mK2*]; *atgls9*[*Punc-25::SOD1(A4V)::YFP*] | DNA6007 |
| *atgEx254*[*Punc-25::GA50::GFP*] | DNA254 |
| *atgEx344*[*Punc-129::SOD1(A4V)::YFP*] | DNA344 |
| *atgEx343*[*Punc-129::SOD1(WT)::YFP*] | DNA343 |
| *atgEx344*[*Pacr-15::SOD1(A4V)::YFP*] | DNA30 |
| *atgEx553*[*Punc-25:: SOD1(A4V)::YFP+ Punc-25::PA50:: mK2*] | DNA553 |
| *atgEx100*[*Pdpy-7::mK2::PH+Pmyo-3::BFP*]; *atgls9*[*Punc-25::SOD1(A4V)::YFP*] | DNA6009 |
| *atgEx60*[*Punc-25::EMTB::mK2*]; *atgls9*[*Punc-25::SOD1(A4V)::YFP*] | DNA6008 |
| *atgls71*[*Punc-25::SNB-1::mK2*]; *atgls9*[*Punc-25::SOD1(A4V)::YFP*] | DNA6010 |
| *atgls320*[*Punc-129:: jRCaMP1b*]; *atgEx10*[*Punc-129::SOD1(A4V)::YFP*] | DNA6015 |
| *atgEx746*[*Punc-25::SOD1(A4V)::PA-GFP*]; *atgls82*[*Punc-25:: RPN-5::mK2*] | DNA746 |
| *atgEx694*[*Punc-25:: SOD1(A4V)::YFP+ Punc25::LMP-1::mK2*] | DNA694 |
| *atgEx601[Punc-25::mK2::RAB-7]; atgIs9[Punc-25::SOD1(A4V)::YFP]* | DNA6017 |
| *atgIs110[Punc-25::mK2::LGG-1]; atgIs9[Punc-25::SOD1(A4V)::YFP]* | DNA6018 |
| *atgEx683[Punc-25:: SOD1(A4V)::YFP::UBQ]* | DNA683 |
| *atgEx647[Punc-25::PA50::GFP::UBQ]* | DNA647 |
| *atgEx626[Punc-25::mK2::SMO-1]; atgIs9[Punc-25::SOD1(A4V)::YFP]* | DNA6026 |
| *atgEx684[Punc-25:: SOD1(A4V)::YFP::SMO-1]* | DNA684 |
| *atgEx869[Punc-25::SOD1(A4V)::FP11+Punc-25::CFP(1-10)::UBQ+ Punc-25::YFP(1-10)::SMO-1]* | DNA869 |
| *atgEx856[Punc-25::GA50::GFP::SMO-1]* | DNA856 |
| *ubc-9(ne4446[ubc-9(G56R)])* | DNA6030 |
| *ubc-9(ne4446[ubc-9(G56R)]); atgIs9[Punc-25::SOD1(A4V)::YFP]* | DNA6031 |
| *ubc-9(ne4446[ubc-9(G56R)]); atgIs20[Punc-25::SOD1(WT)::YFP]* | DNA6032 |

**Supplemental table S2:**
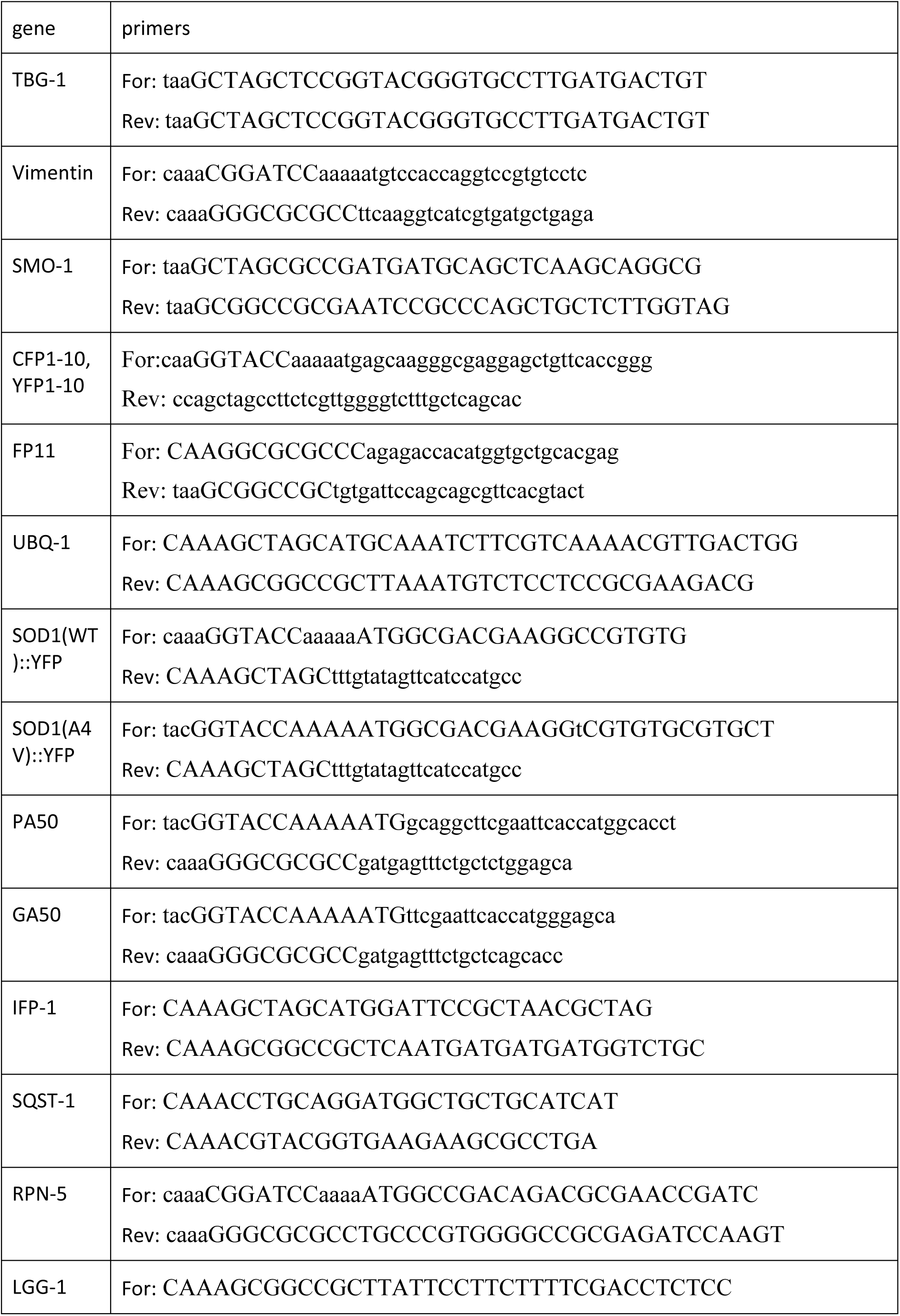
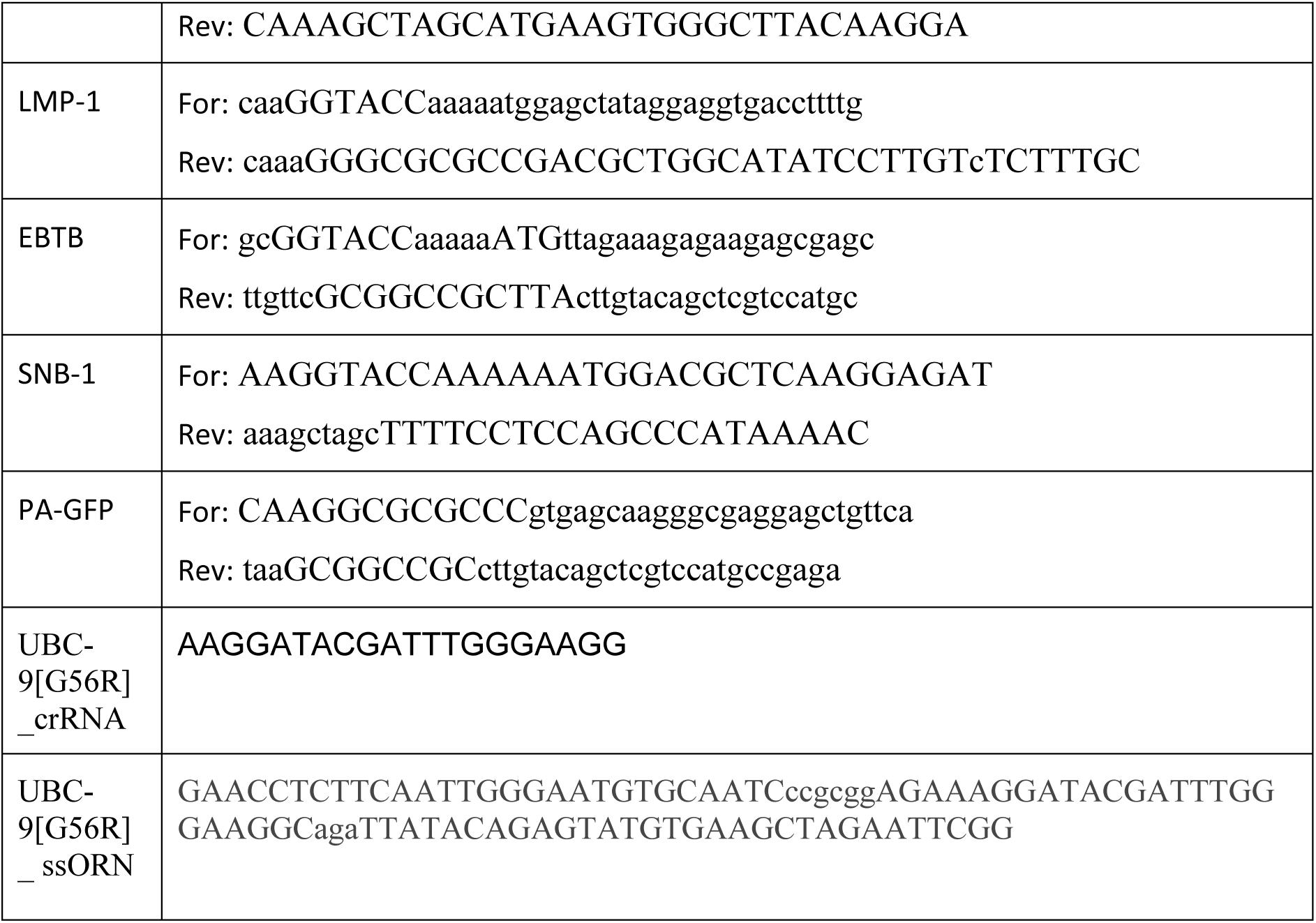
Primer List.

